# Structural basis of flagellar rod assembly on the FliPQR protein-export channel

**DOI:** 10.1101/2025.02.07.637169

**Authors:** Miki Kinoshita, Tomoko Miyata, Fumiaki Makino, Katsumi Imada, Keiichi Namba, Tohru Minamino

## Abstract

The FliPQR complex constitutes a channel for export of the bacterial flagellar proteins involved in axial structure assembly. It also serves as a template for flagellar rod assembly. A periplasmic gate, formed by the N-terminal α-helices of FliP and FliR, remains closed until FliE assembles onto FliP and FliR. The mechanism by which FliE opens the gate and assembles has remained unclear. Here, we present a cryoEM structure of the FliPQR complex in closed form at 3.0 Å resolution. A β-cap formed by the N-terminal β-strands of FliP and FliR creates a tight seal in the closed gate. Interaction of FliE with FliP and FliR induces a conformational change in FliP and FliR, with their N-terminal α-helices move up and outward. Consequently, the N-terminal β-strands of FliP and FliR start opening the periplasmic gate one after another and form a docking site for FliE to initiate rod assembly.

## Introduction

*Salmonella enterica* serovar Typhimurium (hereafter *Salmonella*) employs the flagellar type III secretion system (hereafter referred to as fT3SS) to construct the axial structure of the flagellum, which is responsible for rapid swimming motility in liquid environments^1^. The fT3SS consists of a transmembrane export-gate complex made up of FlhA, FlhB, FliP, FliQ, and FliR and a cytoplasmic ATPase ring complex consisting of FliH, FliI, and FliJ (Fig. 1)^2,3^. The export-gate complex is located within the central pore of the basal body MS-ring and is powered by the transmembrane electrochemical gradient of protons^4,5^. The cytoplasmic ATPase ring complex associates with the C-ring^6^ and acts as an activator that enables the export-gate complex to become a proton-driven protein transporter^7,8^.

**Fig. 1.**
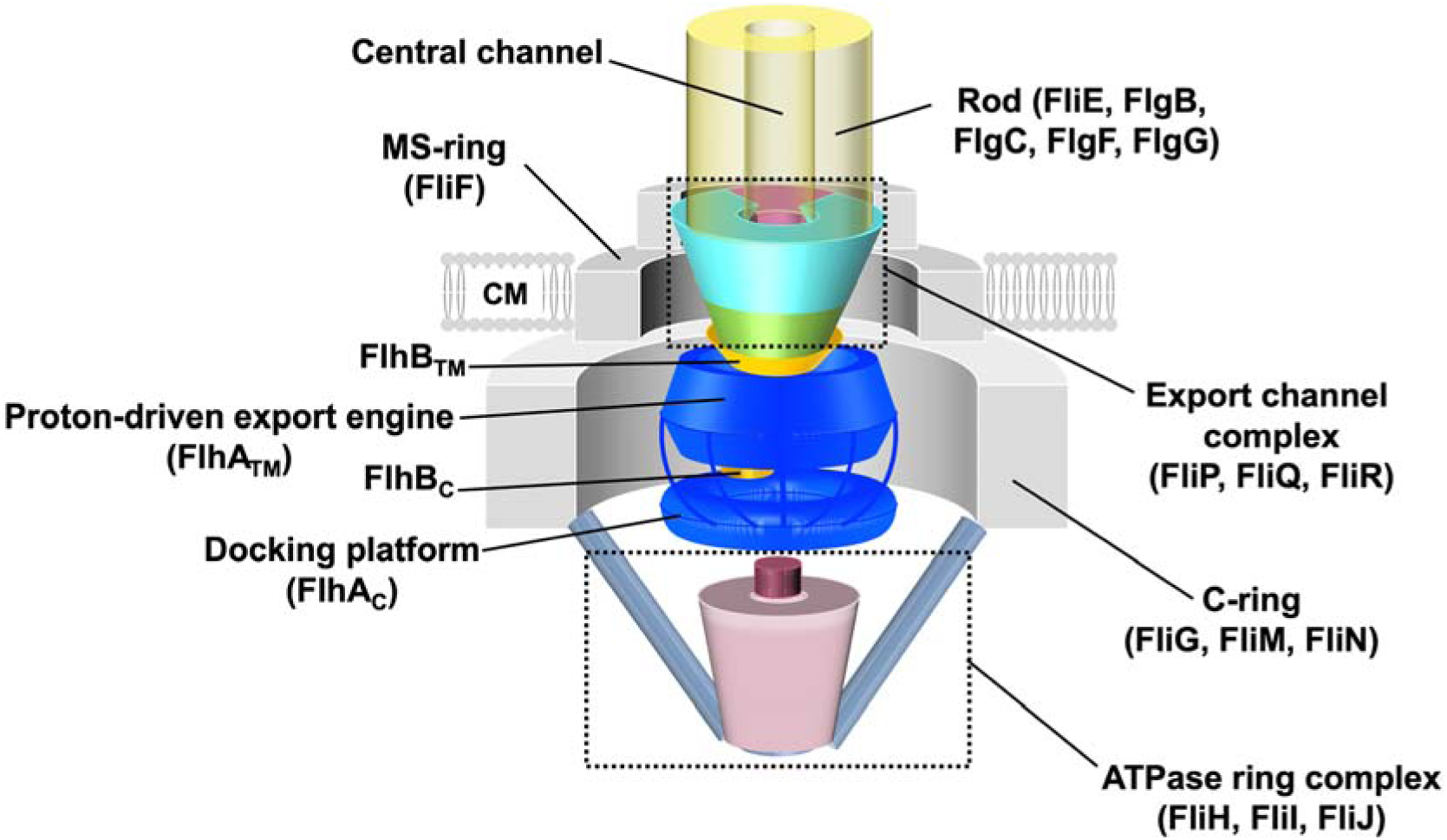
Schematic diagram of the flagellar type III secretion system. The flagellar type III secretion system consists of a transmembrane export-gate complex made up of FlhA, FlhB, FliP, FliQ, and FliR and a cytoplasmic ATPase ring complex consisting of FliH, FliI, and FliJ. The export-gate complex is located within the central pore of the basal body MS-ring, whereas the cytoplasmic ATPase firmly associates with the basal body C-ring through interactions between FliH and FliN. FliP, FliQ, and FliR form a protein export channel for efficient and rapid export of flagellar structural subunits. FlhB associates with the FliPQR complex and plays an important role in opening and closing the cytoplasmic gate of the channel. The N-terminal transmembrane domains of FlhA serve as a proton-driven export engine that couples inward-directed proton flow through the proton channel with outward-directed protein translocation through the export channel. The C-terminal domains of FlhA and FlhB project into the central cavity of the C-ring and act as a substrate docking platform. Because the FliPQR complex has a helical symmetry, FliE and FlgB can directly assemble onto FliP and FliR to form the most proximal part of the rod. CM, cytoplasmic membrane.

Five FliP subunits and one FliR subunit assemble into a right-handed helical structure with a narrow central pore. This pore serves as a channel for export of proteins that constitute several distinct parts of the flagellum^9–11^. Four FliQ subunits peripherally associate on the outside of the FliP_5_-FliR_1_ complex^11–13^. One FlhB subunit associates with the FliP_5_-FliQ_4_-FliR_1_ complex (hereafter FliPQR) to regulate the opening and closing of the cytoplasmic gate of the export channel^14,15^. FlhA assembles to form a homo-nonameric ring with its transmembrane part possibly surrounding the FliPQR-FlhB complex^16,17^ and functions as a proton-driven export engine^18–21^. The C-terminal cytoplasmic domains of FlhA (FlhA_C_) and FlhB (FlhB_C_) project into the cytoplasmic cavity of the C-ring. They form a docking platform for the cytoplasmic ATPase complex, the flagellar export chaperones, and the structural subunits of the flagellum^22–27^. This docking platform ensures the proper order of flagellar protein export during the flagellar assembly process^28–32^.

The flagellar rod is the drive shaft of the flagellar motor, transmitting the motor torque to the hook and filament in the cell exterior to produce thrust for cell swimming. It is a helical tubular structure formed by five different proteins, FliE, FlgB, FlgC, FlgF and FlgG, with approximately 5.5 subunits per turn of the helix. It is composed of six FliE subunits, five FlgB subunits, six FlgC subunits, five FlgF subunits, and twenty-four FlgG subunits^1^. FliE is exported into the periplasm via the fT3SS^33^ and forms the first helical layer of the rod on top of the FliP and FliR subunits of the export-gate complex, followed by the assembly of FlgB, FlgC, FlgF, and FlgG in this order^12,13^. Thus, the FliPQR complex also acts as a template for rod assembly.

FliE has unique features that distinguish it from the other four rod proteins. The *fliE* gene is adjacent to the *fliF* gene that encodes the MS-ring and is located at a different locus from other rod genes that forms the *flgBCDEFGHIJ* operon^1^. Extensive interactions of FliE with FliP and FliR allow other flagellar proteins to efficiently diffuse through the central channel of the growing flagellar structure and assemble at the distal end^34^. FliE has no common structural motifs found in other rod proteins^12,13,35^. These properties suggest that FliE has a somewhat specialized function. Indeed, it has two distinct functional roles in the flagellar assembly process: first as a structural adapter that firmly anchors the rod to the MS-ring and the FliPQR complex, and second, as an export-channel activator of the fT3SS to fully open the export channel^36–38^. FliE is composed of three α-helices: α1, α2, and α3. Helix α1 of FliE binds to the inner wall of the MS-ring, whereas helices α2 and α3 form domain D0, which is the inner core domain of the flagellar axial structure. This D0 domain interacts with FliP, FliR, FlgB, and FlgC within the MS-ring^12,13^. Thus, FliE appears to couple the completion of export channel formation with the initiation of rod assembly. However, it remains how it carries out this function.

In the structure of the purified FliPQR complex (PDB ID: 6R69)^39^, the periplasmic gate of the export channel is closed by intermolecular interactions between the N-terminal α-helices of five FliP subunits and one FliR subunit. In the native basal body (PDB ID: 8WKK), however, the periplasmic gate is held open because of direct binding of six FliE subunits to these α-helices^40^. Unfortunately, the N-terminal region of FliP has not been visualized in either the purified FliPQR complex or the native basal body^11–13,39,40^. As a result, the mechanism by which the closed periplasmic gate of the export channel opens upon FliE assembly has remained obscure.

To address this issue, we conducted cryo-electron microscopy (cryoEM) structural analysis of the purified *Salmonella* FliPQR complex reconstituted in a peptidisc, which is a sheet of short amphipathic bi-helical peptides, and obtained the structure of the completely closed form at 3.0 Å resolution. We also conducted structure-based mutational analyses. We show that the periplasmic gate of the export channel is entirely sealed by the β-cap formed by the N-terminal β-strands of FliP and FliR. We also demonstrate that the interaction of FliE with FliP induces a conformational change in the MTSF motif of FliP (residues 61–64) that results in the outward movement of its N-terminal α-helix. This process sequentially detaches the β-strands from the β-cap to create the next FliE assembly site. Deletion analyses of residues 155–166 of FliP, which form a structural motif called p-loop on the surface of the FliPQR complex facing the inner wall of the MS-ring, demonstrated that the interaction of the p-loop with the MS-ring inner wall stabilizes the open conformation of its N-terminal α-helix. Based on these observations, we propose that the β-cap not only seals the periplasmic gate to limit uncontrolled protein secretion into the periplasm but also functions as a scaffold upon which the newly transported FliE subunits can efficiently assemble.

## Results

### CryoEM structural analysis of the FliPQR complex reconstituted in a peptidisc

FliP has a cleavable signal peptide (residues 1–21) at its N-terminus. The signal peptide is cleaved during membrane insertion^41^. Residues of 22–42 of mature FliP and residues 1–5 of FliR are not visible in the FliPQR complex solubilized with lauryl maltose neopentyl glycol (LMNG) (PDB ID: 6R69) (Fig. 2a,b)^39^. Because detergents often negatively affect protein structure and function, we decided to prepare the FliPQR complex in a detergent-free solution for structural analysis^42,43^. To efficiently and rapidly purify the FliPQR complex, a 10-residue His-tag was added to the C-terminus of FliR. FliP, FliQ, and FliR-His were expressed from a pTrc99-based plasmid in a *Salmonella* strain SJW1368, in which the flagellar *flhDC* master operon required for the expression of all flagellar genes is deleted. Crude membranes were isolated by ultracentrifugation and solubilized by 1% (w/v) LMNG. The His-tagged FliPQR complex was purified by nickel affinity chromatography, followed by size exclusion chromatography. Then, we reconstituted the FliPQR complex in a peptidisc formed by short amphipathic bi-helical peptides^42^. Multiple copies of the peptide wrap around the transmembrane portion of the FliPQR complex to shield the hydrophobic surface, thereby making the complex very soluble in detergent-free solutions. This preparation yielded homogenously dispersed particles readily visualized by cryoEM (Supplementary Fig. 1a).

**Fig. 2.**
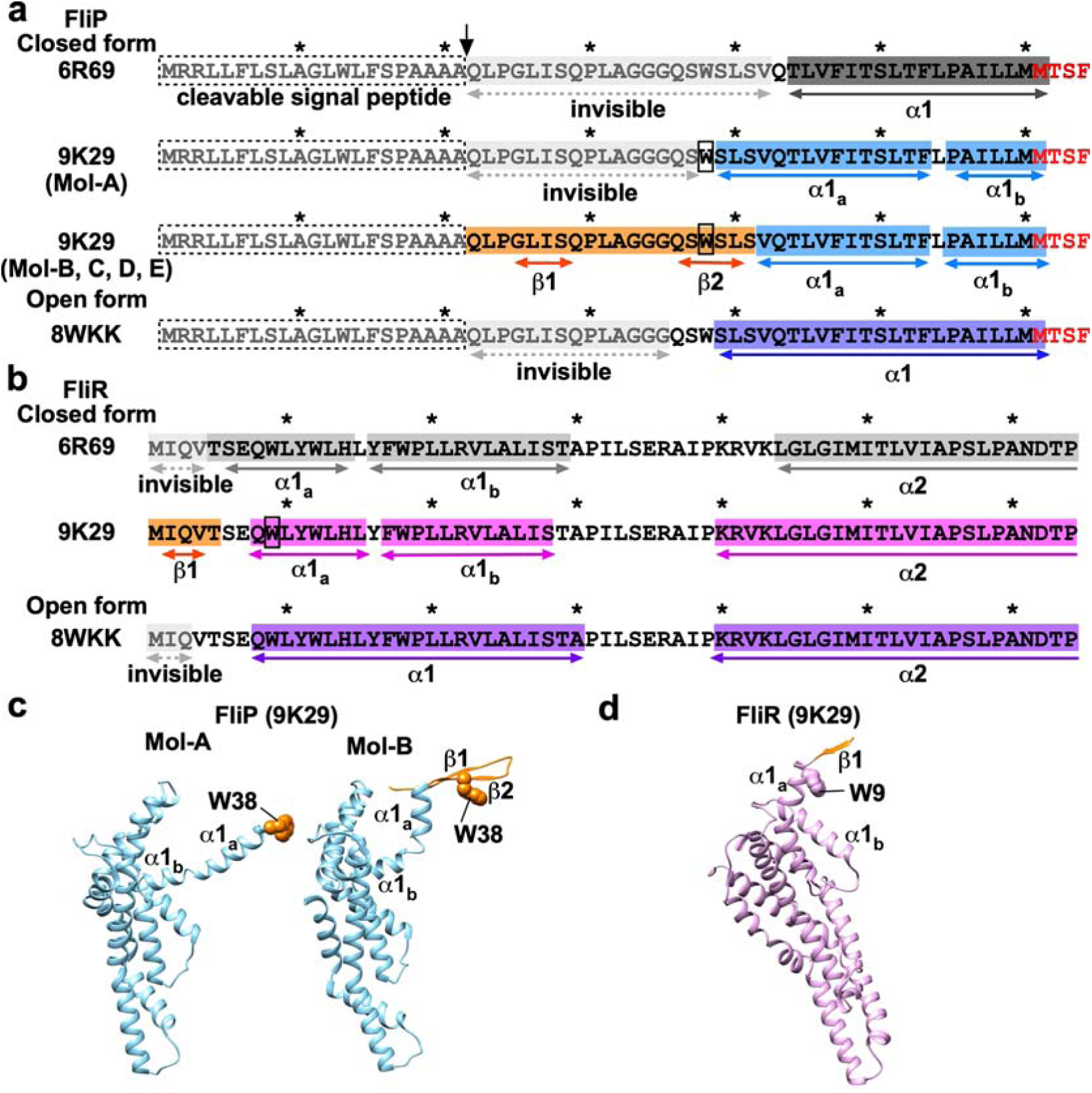
The atomic models of the N-terminal regions of FliP and FliR in a completely closed form of the FliPQR complex. **(a, b)** Primary sequences of the N-terminal regions of (a) FliP and (b) FliR. The N-terminal regions of FliP and FliR are in the closed conformation in the 6R69 and 9K29 structures, but in the open conformation in the 8WKK structure. The first 21 amino acid residues of FliP serve as an N-terminal signal peptide that is cleaved during membrane insertion. The arrow indicates that the cleavage occurs between Ala-21 and Gln-22. Residues 22–42 in the 6R69 structure and residues 22–35 in the 8WKK structure are invisible. However, these residues are visible in the 9K29 structure. The well-conserved MSTF motif is highlighted in red. The first 5 and 3 residues of FliR are not visible in the 6R69 and 8WKK structures, respectively, but are visible in the 9K29 structure. These visible regions, highlighted in orange, form the β-cap, which is stabilized by the six tryptophan residues (Trp-38 of FliP and Trp-9 of FliR) indicated by boxes. **(c)** Cα ribbon diagrams of the atomic models of the two FliP subunits, Mol-A and Mol-B, obtained in this study (PDB ID: 9K29). **(d)** Cα ribbon diagram of the atomic model of FliR obtained in this study (PDB ID: 9K29).

A total of 1,505,398 particles were extracted from 13,230 cryoEM micrographs and analyzed by single-particle image analysis (Supplementary Fig. 1b). After iterative 3D refinement with C1 symmetry, the 3D image of the FliPQR complex was reconstructed at 3.0 Å resolution from 109,333 particles (EMDB ID: EMD-61993) (Supplementary Fig. 1c and Table 1). The map resolution was much better than any of the previous ones^12–14,39,40^. This high-resolution map allowed us to build a more accurate atomic model of the FliPQR complex (PDB ID: 9K29). In agreement with previous reports^12–14,39,40^, the FliPQR complex adopts a right-handed helical structure composed of 5 FliP subunits, 4 FliQ subunits, and 1 FliR subunit (Fig. 3a).

**Table 1.**
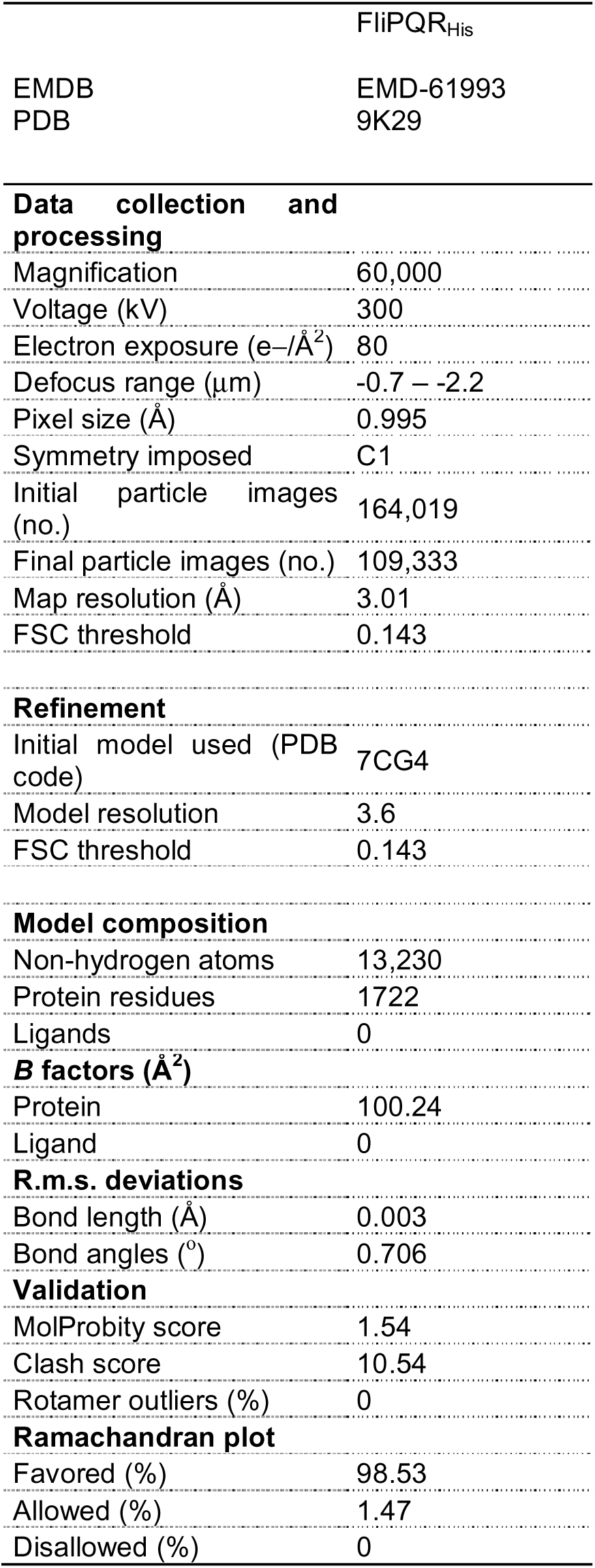
CryoEM data collection, processing, and refinement statistics.

**Fig. 3.**
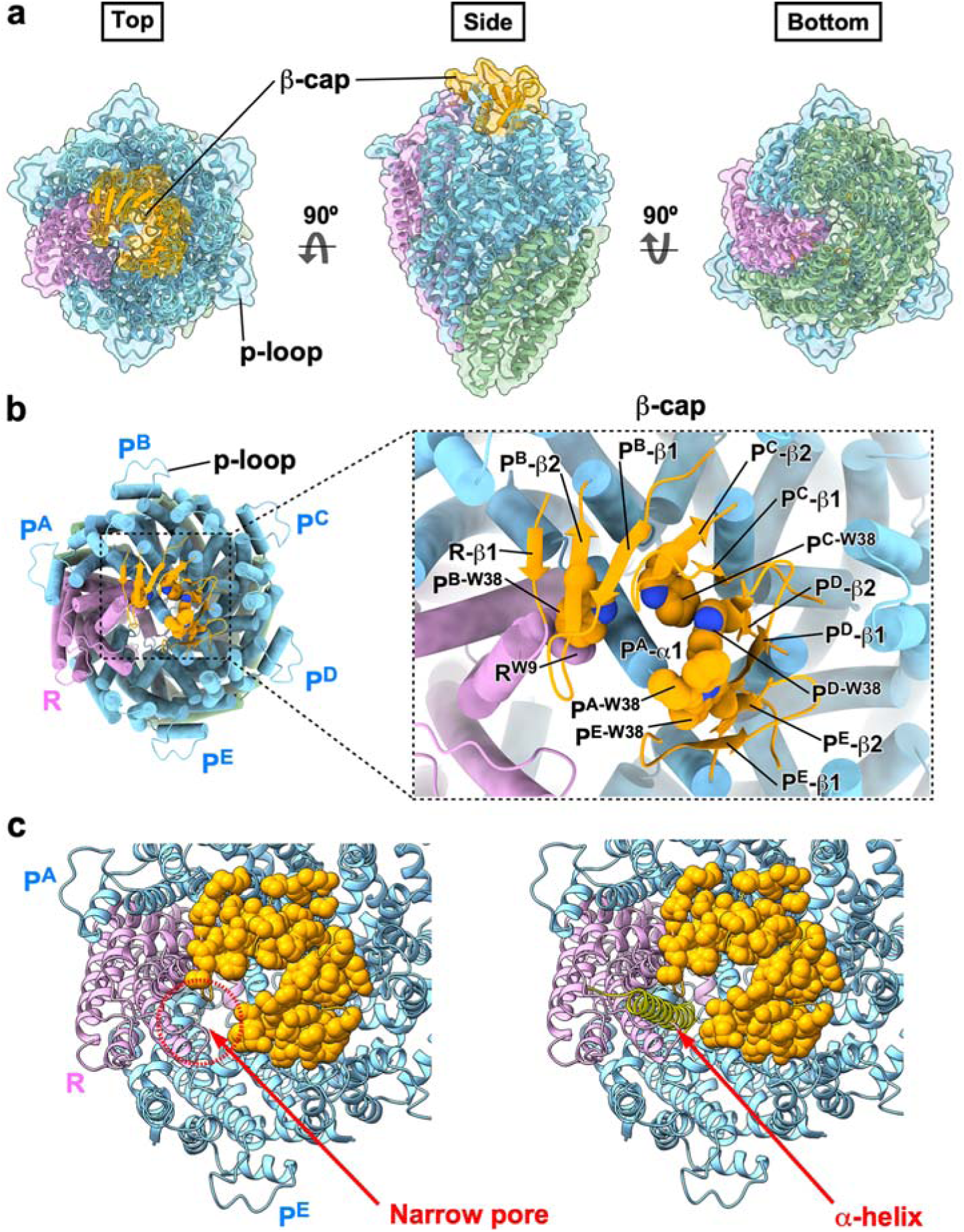
The cryoEM structure of the FliPQR complex reconstituted in the peptidisc. **(a)** Cα ribbon diagram of the atomic model of the FliPQR complex with a stoichiometry of 5 FliP subunits, 4 FliQ subunits, and 1 FliR subunit (PDB ID: 9K29). FliP, FliQ, and FliR are colored in sky blue, dark sea green, and plum, respectively. The newly identified structural elements of FliP and FliR, which are coloured in orange, form the β-cap. The p-loop, consisting of residues 155–166, is located in the outermost part of the FliPQR complex. **(b)** The β-cap formed by the N-terminal β strands of FliP and FliR. Four FliP subunits [Mol-B (P^B^), Mol-C (P^C^), Mol-D (P^D^), and Mol-E (P^E^)], but not the Mol-A (P^A^) subunit, contain two β-strands (β1, β2) at their N-terminal regions, which form a β-hairpin. FliR possesses a β-strand (β1) in its extreme N-terminal region. The four β-hairpins and β1 of FliR form the β-cap, which is stabilized by hydrophobic interactions among six tryptophan residues (Trp-38 of FliP and Trp-9 of FliR). The extreme N-terminal region of α1 of FliP^A^ project into the cavity of the β-cap. **(c)** The narrow pore within the β-cap. The N-terminal region of the Mol-A does not join the β-hairpin structure, which leaves a narrow pore between the FliR and FliP-E subunits in the 9K29 structure (left panel). When the 9K29 structure is superimposed on the equivalent coordinates of the 8WKK structure, it is evident that the pore is sufficiently wide to permit the passage of one α-helix (right panel). Side-chain atoms of amino acid residues of FliP and FliR involved in the formation of the β-cap are indicated by orange spheres.

The densities corresponding to residues 22–41 of four mature FliP subunits, Mol-B (FliP^B^), Mol-C (FliP^C^), Mol-D (FliP^D^), and Mol-E (FliP^E^), and residues 1–5 of FliR were clearly visible, enabling us to build the atomic model of these N-terminal regions (Fig. 2c,d). The density corresponding to residues 22–37 of the Mol-A subunit (FliP^A^) was invisible (Fig. 2a,c). The N-terminal region of the four FliP subunits in which the N-termini are resolved forms a β-hairpin with β-strands β1 and β2. The N-terminal region of FliR contains a single β-strand (β1). These β-strands form a β-sheet at the tip of the FliPQR complex, which we named the β-cap, (Fig. 3). A total of six tryptophan residues, of which five are Trp-38 of FliP and the remaining one is Trp-9 of FliR, stabilize the β-cap structure via hydrophobic contacts with each other (Fig. 3b). Furthermore, the N-terminal α-helix (α1) of FliP^A^ projects toward the export channel, and its N-terminal region, including Trp-38, inserts into the cavity of the β-cap, thereby maintaining the closed state of the periplasmic gate. Consequently, our cyroEM structure of the FliPQR complex adopts a bud-like shape (Fig. 3a, middle panel).

Helix α1 of FliP^A^ is longer by three residues (S39-L40-S41) than helix α1 of the remaining four FliP subunits (Fig. 2a), indicating that the S39-L40-S41 region is convertible between α-helix and β-strand. Helix α1 is kinked at Leu-54 and is divided into two distinct parts, designated as α1_a_ and α1_b_ (Fig. 2a,c). The five FliP subunits are arranged in a helical array, with FliP^A^ at the top and FliP^B^, FliP^C^, FliP^D^, and FliP^E^ descending in a spiral staircase. Except for FliP^A^, α1_a_ rises upward and move outward. In addition, the β-hairpins of the FliP^B^, FliP^C^, FliP^D^, and FliP^E^ subunits also rise upward (Fig. 4a, middle panel), enabling them to form the β-cap together with β1 of FliR (Fig. 3b).

**Fig. 4.**
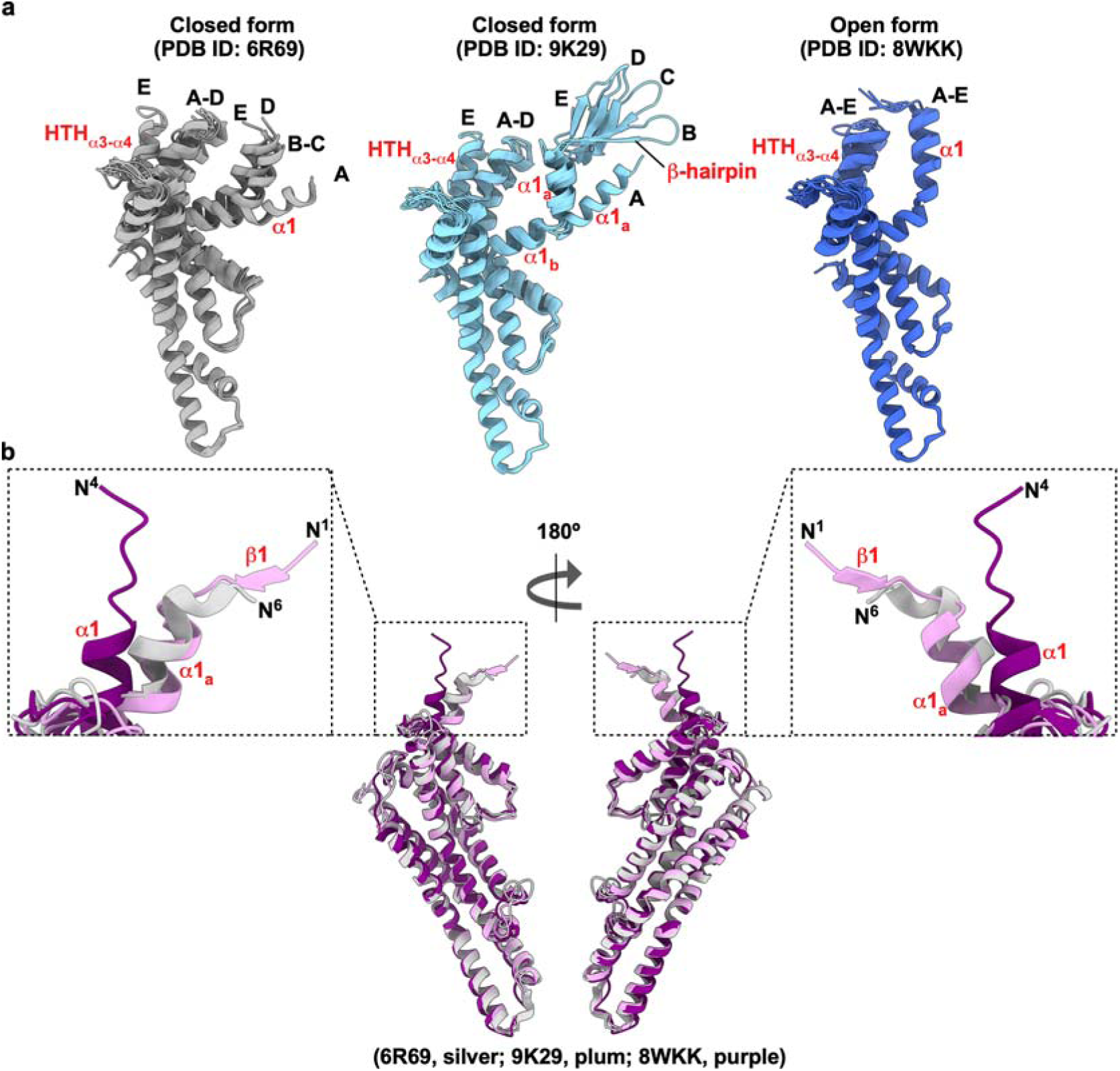
Structural comparisons of the FliPQR complex obtained in this study (PDB ID: 9K29) with the equivalent coordinates of the 6R69 and 8WKK structures. **(a)** Structural comparisons of the five FliP subunits in the three different FliPQR structures; the conformations of the five FliP subunits are slightly different from each other in the 6R69 (dark grey) and 9K29 (sky blue) structures, whereas the five FliP subunits are nearly identical to each other in the 8WKK structure (royal blue). In the 9K29 structure, the N-terminal β-hairpin structure is visible in Mol-B, Mol-C, Mol-D, and Mol-E, but not in Mol-A. The orientation of α1 is different among the five FliP molecules in the 6R69 and 9K29 structures. In the Mol-E subunit, the helix-turn-helix structure (HTH_α3-α4_), which is formed by α3 and α4, and a loop connecting these two helices, is shifted outward relative to the other four FliP subunits. In the 8WKK structure, the orientation of α1 and the position of HTH_α3-α4_ are nearly identical among the five FliP molecules. **(b)** Structural comparison of the FliR subunits in the three different FliPQR structures. The overall structure of FliR is almost identical among the 6R69 (silver), 9K29 (plum), and 8WKK (purple) structures, but in the 8WKK structure, the N-terminal α1 helix is oriented differently than in the 6R69 and 9K29 structures. The N-terminal β-strand (β1) is visible only in the 9K29 structure.

The β-cap has a narrow pore between FliP^E^ and FliR that is wide enough to allow the passage of a single α-helix (Fig. 3c). The FliE-Bla fusion protein is secreted via the fT3SS into the periplasm to a significant degree even in the absence of FliE, suggesting that FliE can pass through the pore^38^. When the corresponding atoms of the 9K29 and 8WKK structures are superimposed, it is very clear that the pore can properly and precisely accommodate helix α3 of the first FliE subunit (FliE^1^) at the first FliE assembly site (Fig. 3c, right panel). Therefore, we propose that the β-cap not only keeps the periplasmic gate closed until FliE assembles onto FliP and FliR but also serves as a scaffold for the initiation of FliE assembly.

### Structural comparison of the open and closed forms of the FliPQR complex

The periplasmic gate of the export channel is closed in the 6R69 and 9K29 structures, but it is open in the 8WKK structure (Supplementary Fig. 2). To investigate how the β-cap disassembles when the periplasmic gate adopts the open conformation, we superimposed the FliP, FliQ, and FliR subunits in the 6R69 and 8WKK structures on the corresponding ones obtained in this study (PDB ID: 9K29). The overall structures of FliP, FliQ, and FliR in the 9K29 structure are nearly identical to those in the 6R69 and 8WKK structures except for the conformation and orientation of the N-terminal regions of FliP and FliR (Fig. 4 and Supplementary Figs. 3 and 4).

Helix α1 of all five FliP subunits in the 6R69 structure is five residues shorter than helix α1 of FliP^A^ in the 9K29 structure and two residues shorter than those observed in the remaining four FliP subunits (Fig. 2a). Furthermore, the first four residues of FliR are invisible in the 6R69 structure (Fig. 2b). These observations suggest that the β-cap stabilizes the intermolecular FliP-FliP and FliP-FliR interactions, which in turn ensures that α1 is maintained in its closed conformation. In the 8WKK structure, the length of α1 observed for all five FliP subunits is the same as that of FliP^A^ in the 9K29 structure. In the native basal body, α1 of FliP interacts with FliE to stabilize the extreme N-terminal region of α1 (Supplementary Fig. 5).

The orientation of α1 is different among the five FliP molecules in the closed structures of 6R69 and 9K29. In the FliP^E^ subunit, the helix-turn-helix structure (HTH_α3-α4_), which is formed by α3 and α4, and a loop connecting these two helices, is shifted outward relative to the other four FliP subunits (Fig. 4a, left and middle panels). Conversely, in the open structure of 8WKK, the orientation of α1 and the position of HTH_α3-α4_ are nearly identical among the five FliP molecules (Fig. 4a, right panel). When each of the five FliP subunits in the 6R69 and 8WKK structures is superimposed on the corresponding FliP subunit in the 9K29 structure, α1 rises upward and HTH_α3-α4_ moves outward in the 8WKK structure (Supplementary Fig. 3).

The I2-Q3-V4 sequence of FliR forms strand β1 in the 9K29 structure, whereas these three residues are invisible in the 8WKK structure (Fig. 2b). This indicates that β1 is stable only in the β-cap. Helix α1 of FliR is kinked at Tyr-16 in the 9K29 structure, thereby dividing it into two distinct parts, named α1_a_ and α1_b_ (Fig. 2d). This kink allows β1 to form an antiparallel β-sheet with β2 of FliP^B^ in the β-cap. Additionally, it enables α1_a_, including Trp-9, to orient toward the export channel to stabilize the β-cap (Fig. 3b). In the 8WKK structure, α1 of FliR moves outward as this kink disappears (Fig. 4b).

The extensive hydrophobic interactions of FliE and FlgB with FliP and FliR firmly fix the rod onto the FliPQR complex^12,13,40^. The hook-capping protein FlgD and the hook protein FlgE are secreted into the periplasm in the *flgB* mutant, but the *fliE* mutation severely inhibits the secretion of these two proteins^34,36,37^. Therefore, we suggest that the extensive hydrophobic interactions of FliE with the FliP and FliR subunits not only disrupt the β-cap but also induce conformational changes in α1 of FliP and FliR to fully open the periplasmic gate of the export channel.

### Rationale for mutational analyses

Until FliE binds to FliP and FliR, the periplasmic gate of the export channel resembles a closed flower bud because of the β-cap at the tip of the FliPQR complex. When FliE interacts with FliP and FliR, the periplasmic gate opens completely, resembling a flower in full bloom. Comparison of the open and closed conformations of FliP shows that, although the conformational changes between the open and closed structures differ slightly from subunit to subunit (Fig. 4a), we found a consistent conformational change in the region of residues 59–64 in all five subunits. The well-conserved MTSF motif is located in this region, rendering it an appropriate subject for mutational analysis (Fig. 5a). The central question addressed is whether the MTSF motif plays a crucial role in the function of the periplasmic gate. Networks of hydrophobic side-chain interactions surrounding the MTSF motif are responsible for maintaining the open and closed conformations of the MTSF motif. Therefore, the second group of candidates for mutational analysis comprises residues that constitute the side-chain interaction networks surrounding the MTSF motif (Fig. 6 and Supplementary Fig. 6).

**Fig. 5.**
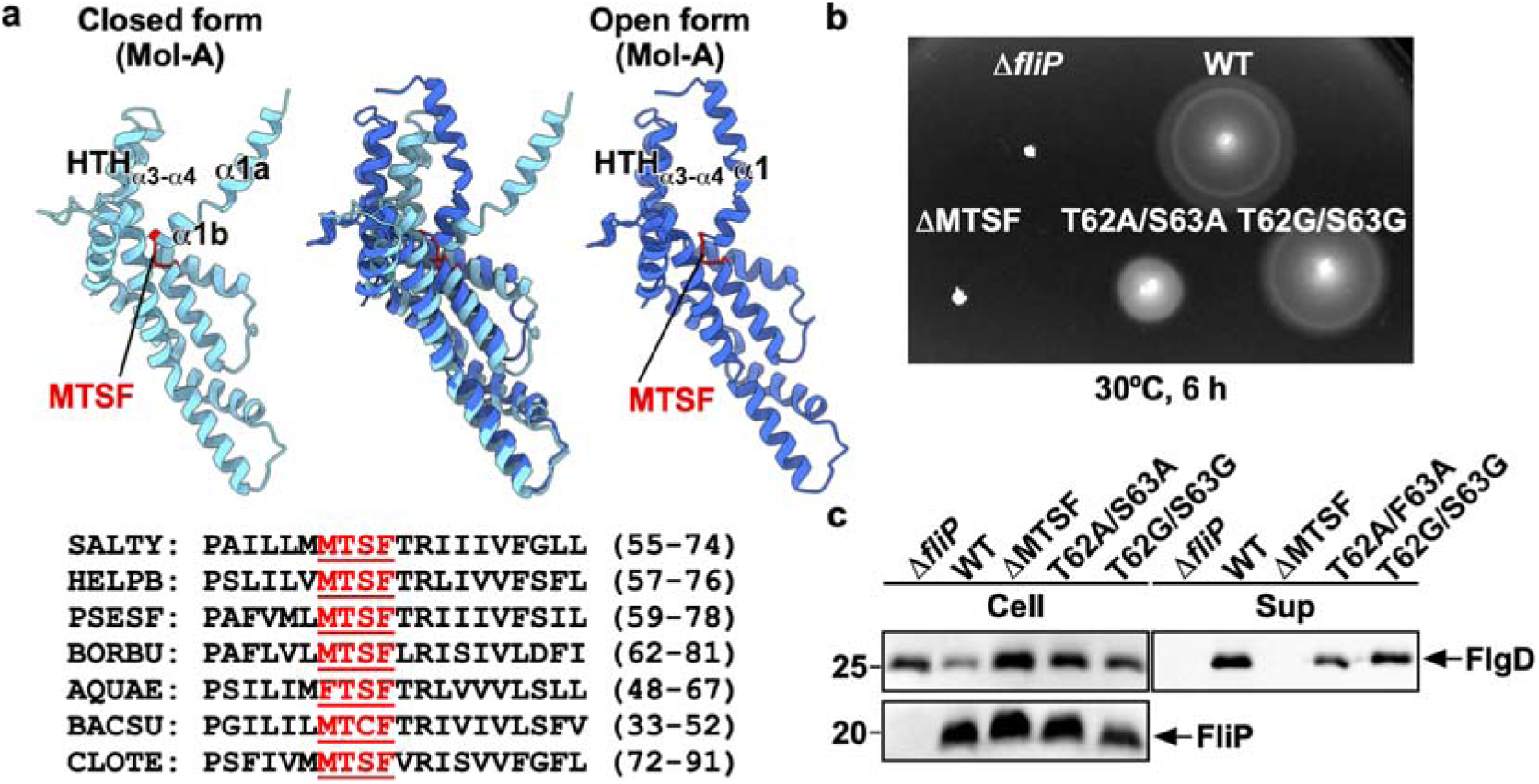
Mutational analysis of the conserved MTSF motif in FliP. **(a)** Structural comparison and multiple sequence alignments of the conserved MTSF motif of FliP. The closed (PDB ID: 9K29, sky blue) and open (PDB ID: 8WKK, royal blue) forms of the Mol-A subunit are shown. Multiple sequence alignment was carried out by Clustal Omega. The highly conserved MTSF motif of FliP is highlighted in red. UniProt Accession numbers: *Salmonella enterica* (SALTY), C7BZE6; *Helicobacter pylori* (HELPB), C7BZE6; *Pseudomonas syringae* (PSESF), A0A2V0QEJ7; *Borreliella burgdorferi* (BORBU), Q44763; *Aquifex aeolicus* (AQUAE), O67750; *Bacillus subtilis* (BACSU), P35528; *Clostridium tetani* (CLOTE), Q893Z3. **(b)** Motility of a *Salmonella fliP* null mutant harbouring pTrc99AFF4 (indicated as Δ*fliP*), pKY69 (indicated as WT), pMKM69(ΔMTSF) (indicated as ΔMTSF), pMKM69(T62A/S63A) (indicated as T62A/S63A), and pMKM69(T62G/S63G) (indicated as T62G/S63G) in soft agar. The plate was incubated at 30°C for 6 hours. At least seven independent assays were performed, **(c)** Flagellar protein secretion assays. Whole cell proteins (Cell) and culture supernatant fractions (Sup) were prepared from the above transformants. A 5 μl solution of each protein sample, which was normalized to an optical density of OD_600_, was subjected to SDS-PAGE, followed by immunoblotting with polyclonal anti-FlgD (first row) or anti-FliP (second row) antibody. The positions of molecular mass markers (kDa) are shown on the left. At least three independent assays were carried out.

**Fig. 6.**
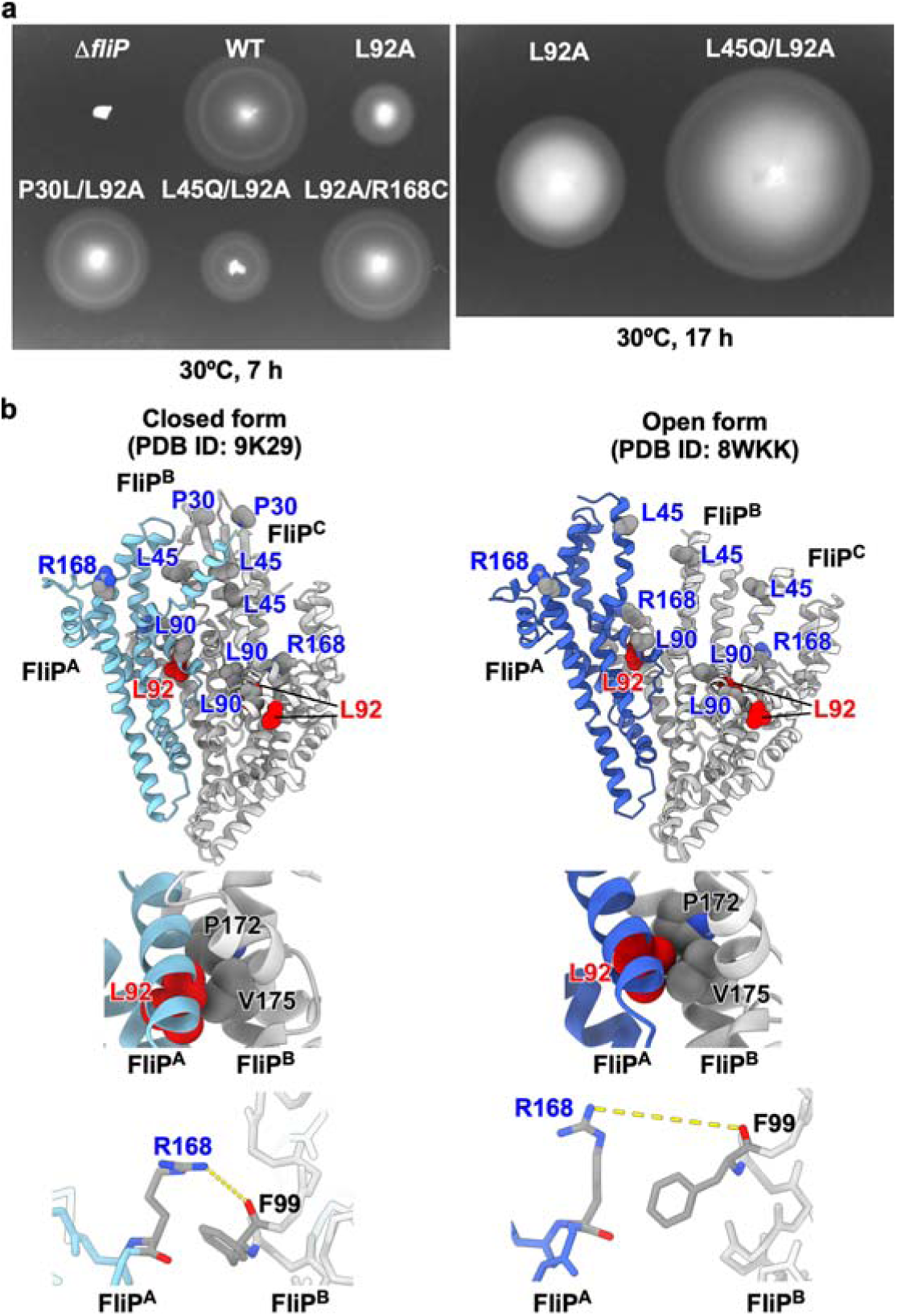
Isolation of gain-of-function mutants from the *fliP(L92A)* mutant. **(a)** Motility of a *Salmonella fliP* null mutant carrying pTrc99AFF4 (indicated as Δ*fliP*), pKY69 (indicated as WT), pMKM69(L92A) (indicated as L92A), pMKM69(L92A)-SP1 (indicated as P30L/L92A), pMKM69(L92A)-SP2 (indicated as L45Q/L92A), and pMKM69(L92A)-SP3 (indicated as L92A/R168C) in soft agar. The plate was incubated at 30°C. At least seven independent assays were carried out. **(b)** Location of intragenic suppressor mutation sites isolated from the *fliP(L92A)* mutant. Three FliP subunits [Mol-A (FliP^A^), Mol-B (FliP^B^), and Mol-C (FliP^C^)] are shown. The L92A mutation site and its intragenic suppressor mutation sites are highlighted in red and blue, respectively. Leu-92 of Mol-A makes hydrophobic contacts with both Pro-172 and Val-175 of Mol-B in the open form of FliP, but only with Val-175 in the closed form of FliP. Pro-30 is located within the β-cap in the closed structure. The conserved Leu-45 and Leu-90 residues are involved in the FliP-FliE interaction in the open structure (see Supplementary Fig. 5). Arg-168 of Mol-A forms a hydrogen bond with the carbonyl group of Phe-99 of Mol-B in the closed structure but not in the open structure.

Another mutational analysis was designed to elucidate the mechanism by which the FliE-FliP interaction induces remodelling of the hydrophobic interaction networks to enable FliP to adopt the open conformation. The region comprising residues 155–166 of FliP is designated as the p-loop and is located in the outermost part of the FliPQR complex (Fig. 3). This p-loop gets closer to the i-loop of FliF in inner wall of the MS-ring when FliP adopts the open conformation (Fig. 7a). Consequently, the p-loop is a prime candidate for mutational analysis to determine its role in stabilizing the open conformation of the FliPQR complex inside the MS-ring.

**Fig. 7.**
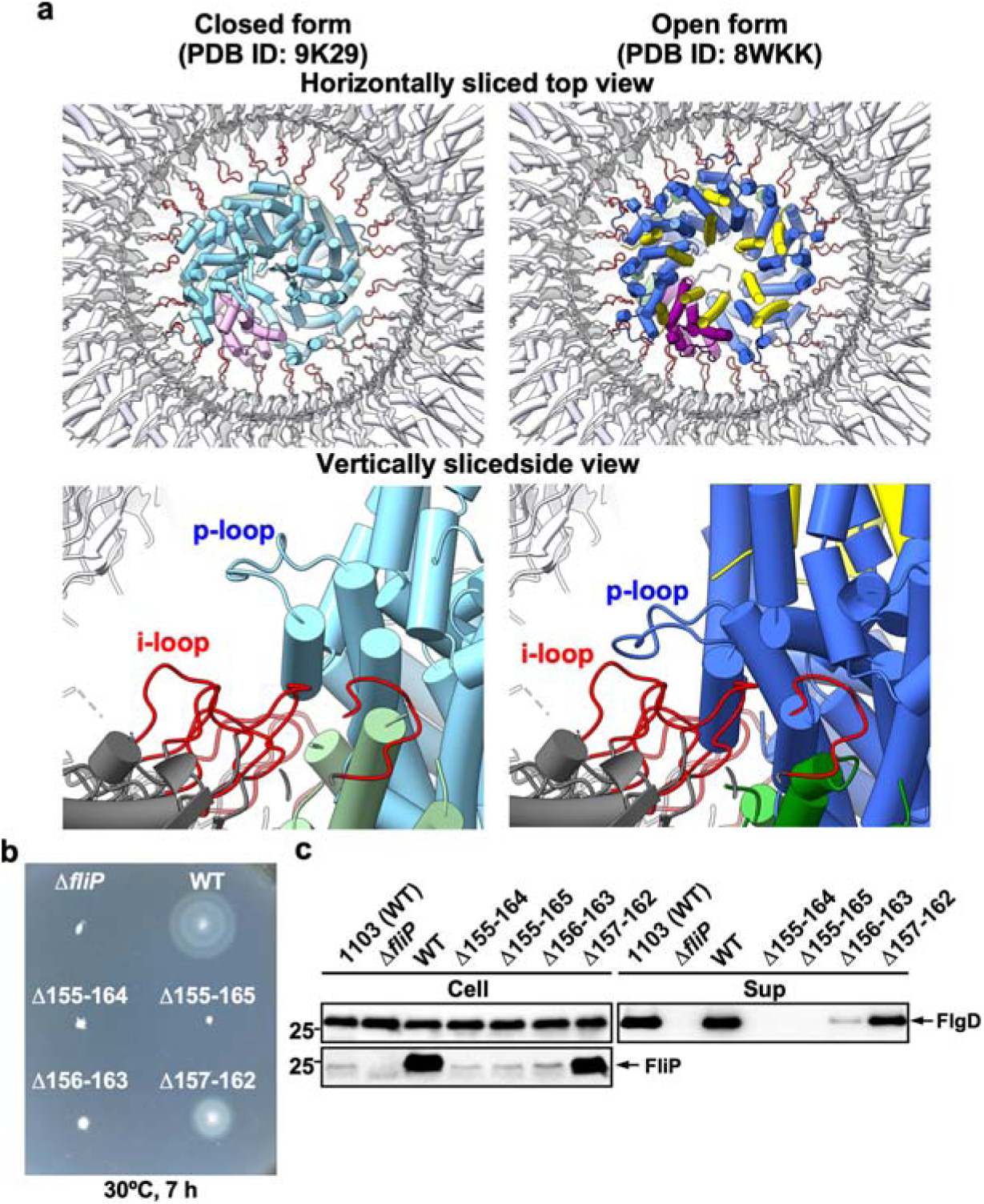
Effect of in-frame deletions of the p-loop of FliP on flagellar protein export. **(a)** Structural comparison of the p-loop conformations in the closed (PDB ID: 9K29) and open (PDB ID: 8WKK) forms of the FliPQR complex. The open and closed structures are superimposed on the equivalent coordinates of the 7NVG structure. FliP, FliQ, and FliR are colored in sky blue, dark sea green, and plum, respectively, in the 9K29 structure. FliE, FliP, FliQ, and FliR are colored in yellow, royal blue, green, and purple, respectively, in the 8WKK structure. The FliPQR complex is located within the central pore of the MS-ring. Each p-loop (residues 155–166) makes physical contact with the i-loop of the MS-ring protein, FliF (red, residues 159 –172 in FliF) within the MS-ring after 6 FliE subunits assemble on FliP and FliR. **(b)** Motility of a *Salmonella fliP* null mutant harbouring pTrc99AFF4 (indicated as Δ*fliF*), pKY69 (indicated as WT), pMKM69(Δ155-164) (indicated as Δ155-164), pMKM69(Δ155-165) (indicated as Δ155-165), pMKM69(Δ156-163) (indicated as Δ156-163), and pMKM69(Δ157-162) (indicated as Δ157-162),in soft agar. The plate was incubated at 30°C for 7 hours. At least seven independent assays were carried out. **(c)** Flagellar protein secretion assays. Whole cell proteins (Cell) and culture supernatant fractions (Sup) were prepared from the above transformants. A 5 μl solution of each protein sample, which was normalized to an optical density of OD_600_, was subjected to SDS-PAGE, followed by immunoblotting with polyclonal anti-FlgD (first row) or anti-FliP (second row) antibody. The positions of molecular mass markers (kDa) are shown on the left. At least three independent assays were performed.

### Mutational analysis of the MTSF motif of FliP

Met-61, Thr-62, Ser-63, and Phe-64 are highly conserved among FliP homologues. They form a hinge that allows helix α1 of FliP to move upward when the FliPQR complex assumes the open conformation (Fig. 5a). To investigate whether a conformational rearrangement of the MTSF motif is necessary to open the periplasmic gate completely, we generated the *fliP*(ΔMTSF) mutant and analyzed its motility in soft agar (Fig. 5b). Immunoblotting using polyclonal anti-FliP antibody revealed that the deletion did not affect the cellular level of FliP (Fig. 5c). However, unlike wild-type FliP, the ectopic production of FliP(ΔMTSF) did not restore motility in the Δ*fliP* mutant (Fig. 5b).

To test whether the lack of motility is a consequence of defective export of flagellar proteins, we analyzed the secretion of FlgD, a representative export substrate of the fT3SS. Immunoblotting with polyclonal anti-FlgD antibody revealed that the MTSF deletion inhibits FlgD secretion (Fig. 5c). Thus, the MTSF motif of FliP is necessary for the initiation of flagellar protein export.

To test whether the conformational flexibility of the MTSF motif is a prerequisite for the movement of α1, we generated two *fliP* mutants, *fliP*(T62A/S63A) and *fliP*(T62G/S63G). The T62A/S63A double substitution caused a decrease in both motility and FlgD secretion. However, the T62G/S63G double substitution had no impact on either motility or FlgD secretion (Fig. 5b,c). Because these two double mutations did not affect the cellular level of FliP (Fig. 5c), we suggest that the important function of the MTSF motif is to provide the conformational flexibility that allows α1 of FliP to move in a direction that completely opens the periplasmic gate.

### Role of the conserved Leu-92 residue of FliP

As detailed in Supplementary Result 1, the *fliP*(L92A) mutation significantly decreased motility (Fig. 6a). Leu-92, situated at the interface of the FliP subunits, forms hydrophobic interactions with Val-175 of the neighbouring FliP subunit in the closed form and with both Pro-172 and Val-175 of the neighbouring FliP subunit in the open form (Fig. 6b). Thus, Leu-92 may enhance hydrophobic interactions between the FliP subunits in the open structure of the FliPQR complex. The L92A substitution reduces the stability of the open conformation of FliP, thereby increasing the likelihood that helix α1 will adopt a closed conformation.

Differences in the hydrophobic interaction mode of Leu-92 between the open and closed states of the FliPQR complex can be attributed to two factors: the 0.5-turn shift of the α-helix containing Val-175 relative to Leu-92 and the different orientation of the side chain of Leu-92. Therefore, we introduced four *fliP* mutations that cause the residue substitutions, L90A/G91A, L90A/L92A, G91A/L92A, and L90A/G91A/L92A. None of these changes affected the cellular level of FliP (Supplementary Fig. 6b, upper panel). The motility of the *fliP*(L90A/L92A) and *fliP*(L90A/G91A/L92A) mutants was much better than that of the *fliP*(L92A) mutant although not as good as that of the wild type (Supplementary Fig. 6c, left panel). Because the G91A substitution alone did not improve the motility of the *fliP*(L92A) mutant, we conclude that the L90A substitution allows FliP(L92A) to adopt a stable open conformation. In the closed structure of 9K29, Leu-90 establishes an intramolecular hydrophobic contact with Phe-64 in the MTSF motif (Supplementary Fig. 7a). In contrast, Leu-90 interacts with Met-102 of FliE in the open structure (Supplementary Fig. 5, lower panel). Therefore, we suggest that the interaction between FliP and FliE induces a restructuring of the hydrophobic side-chain networks surrounding the MTSF motif, thereby opening the periplasmic gate of the export channel.

We also isolated intragenic suppressor mutants from the *fliP*(L92A) mutants. DNA sequencing revealed that the *fliP*(P30L), *fliP*(L45Q) and *fliP*(R168C) mutations restored the motility of the *fliP*(L92A) mutant (Fig. 6a). Pro-30 is located within the β-cap, which serves to stabilize the closed conformation of the periplasmic gate (Fig. 6b). Leu-45 is situated at the core of the hydrophobic region between the α1 helices of FliP subunits in the closed form of the FliPQR complex (Supplementary Figs 8 and 9). Arg-168 forms a hydrogen bond with the carbonyl group of Phe-99 of the adjacent FliP subunit in the closed form. However, this hydrogen bond is absent in the open form (Fig. 6b). This indicates that Arg-168 plays a role in stabilizing the closed conformation of FliP. Therefore, these three residues stabilize the closed conformation, and the P30L, L45Q, and R168C substitutions destabilize the closed conformation of the periplasmic gate even in the presence of the *fliP*(L92A) mutation.

### Interaction between FliF and FliP

The i-loop, residues 159–172 of FliF, is required for assembly of the export-gate complex in the central pore of the MS-ring^44,45^. Residues 155–165 of FliP (p-loop) engage with the i-loop when FliP adopts an open conformation induced by interaction between FliP and FliE (Fig. 7a, right panels). However, the p-loop does not interact with the i-loop when FliP adopts a closed conformation (Fig. 7a, left panels). This raises the possibility that the interaction between the p-loop of FliP and the i-loop of FliF stabilizes the open conformation of the periplasmic gate.

To test this hypothesis, we constructed four *fliP* deletion mutants, Δ155–164, Δ155–165, Δ156–163, and Δ157–162. Deletion of residues 157–162 decreased both motility (Fig. 7b) and FlgD secretion (Fig. 7c) but, based on immunoblotting, did not decrease the level of FliP. Thus, residues 157–162 are required for efficient flagellar protein export but not for stability of FliP.

The other deletions had more severe effects. The Δ156–163 mutant had markedly diminished motility and reduced FlgD secretion, and this was possibly due to a significant reduction in the cellular FliP level. So, the deletion of two flanking residues 157-162 severely affected the stability of FliP. The larger deletions Δ155– 164 and Δ155–165 resulted in a completely non-motile phenotype (Fig. 7b,c) as well as a significant decrease in the cellular level of FliP (Fig. 7c). However, the FliP levels of these three deletion variants were still comparable to the level of FliP expressed from the chromosomal *fliP* gene. These results show that the intact p-loop is required for FliP to adopt the open conformation upon FliE assembly onto FliP and FliR. The decreased stability of FliP in the Δ155–164, Δ155–165, and Δ156–163 mutants additionally show that some elements of the p-loop are also required for proper folding of FliP.

## Discussion

It has previously been shown that the FliPQR complex provides a channel for the export of proteins that assemble into the rod, the hook, and the filament and that it provides a structural template for rod assembly within the MS-ring^12,13^. The N-terminal α1 helices of FliP and FliR serve as a periplasmic gate of the export channel. The extensive hydrophobic interactions between these helices close the periplasmic gate until FliE assembles onto FliP and FliR to form the first layer of the proximal rod at the tip of the FliPQR complex. The assembly of FliE also opens the periplasmic gate to allow export of the proteins that compose the rod, the hook, and the filament^11–13,38^. Thus, FliE assembly is an important checkpoint for the sequential flagellar assembly process. However, the detailed mechanism for opening the periplasmic gate and creating the export channel was not fully understood.

We therefore performed high-resolution cryoEM image analysis of the FliPQR complex reconstituted in peptidisc. This allowed us to visualize the β-cap at the tip of the FliPQR complex (Fig. 3a). The β-cap is stabilized by hydrophobic interactions between Trp-38 of five FliP subunits and Trp-9 of one FliR subunit. In addition, the extreme N-terminus of α1 of one FliP protein (FliP^A^) projects into the cavity of the β-cap, providing a tight seal for the periplasmic gate (Fig. 3b).

To determine how the interactions of FliP and FliR with FliE disrupt the β-cap and allow the α1 helices of FliP and FliR to adopt the conformation they take in the open gate, we performed mutational analyses. The conserved Leu-92 residue stabilizes the open conformation through interactions with Pro-172 and Val-175 of its neighbouring FliP subunit. The L92A substitution removes these hydrophobic contacts and causes the gate to remain closed much of the time, consequently impairing flagellar protein export and motility significantly (Fig. 6).

In the closed gate, Leu-90 of FliP makes an intramolecular hydrophobic contact with Phe-64 within the MTSF motif (Supplementary Fig. 7a) but establishes a hydrophobic interaction with Met-102 of FliE in the open form (Supplementary Fig. 5, lower panel). Pro-30 is located within the β-hairpin and stabilizes the β-cap in the closed form (Fig. 6b). In the closed form, Leu-45 is in helix α1, where it forms a hydrophobic core with Phe-47 and Leu-51 that keeps the export channel closed, as detailed in Supplementary Result 2 (Supplementary Figs. 8 and 9). In the open form, helix α1 including Leu-45 interacts with FliE (Supplementary Fig. 5, upper panels).

The conformational flexibility of the MTSF motif of FliP is necessary for opening the periplasmic gate (Fig. 5). It seems that the extensive hydrophobic interactions of FliP with FliE induce a remodelling of the hydrophobic side-chain networks surrounding the MTSF motif. These changes allow Leu-92 to interact with both Pro-172 and Val-175 of its neighbouring FliP subunit to stabilize the conformation of the MTSF motif in the open gate (Fig. 6b). This promotes the release of the β-hairpin from the β-cap, and helix α1 of FliP adopts the conformation that it assumes in the open gate. Furthermore, the direct contact of each p-loop with the i-loop of FliF locks the open state of each FliP subunit within the MS-ring (Fig. 7). The second-site substitutions P30L, L45Q, L90A, and R168C destabilize the closed conformation conferred by FliP(L92A), thereby improving the motility of the *fliP*(L92A) mutant to a significant degree (Fig. 6a and Supplementary Fig. 6c, left panel).

How does FliE gain access to the distal face of the FliPQR complex? The β-cap tightly seals the periplasmic gate, but a narrow pore remains open between FliR and FliP^E^ that is wide enough to permit the passage of a single α-helix (Fig. 3c, left panel). Helix α1 of FliP^A^ projects toward the β-hairpin of FliP^E^ in the closed form of the FliPQR complex, and its Trp-38 residue makes a hydrophobic contact with Trp-38 of FliP^E^ (Fig. 3b). As a result, helix α1 of FliP^A^ efficiently and properly directs the N-terminal helix of FliE^1^ into the pore. Once secreted, FliE^1^ initiates the assembly of FliE subunits on the scaffold provided by the β-cap, which then leads to the disruption of the β-cap and the conformational changes in FliP and FliR that generate the open form of the gate (Fig. 8).

**Fig. 8.**
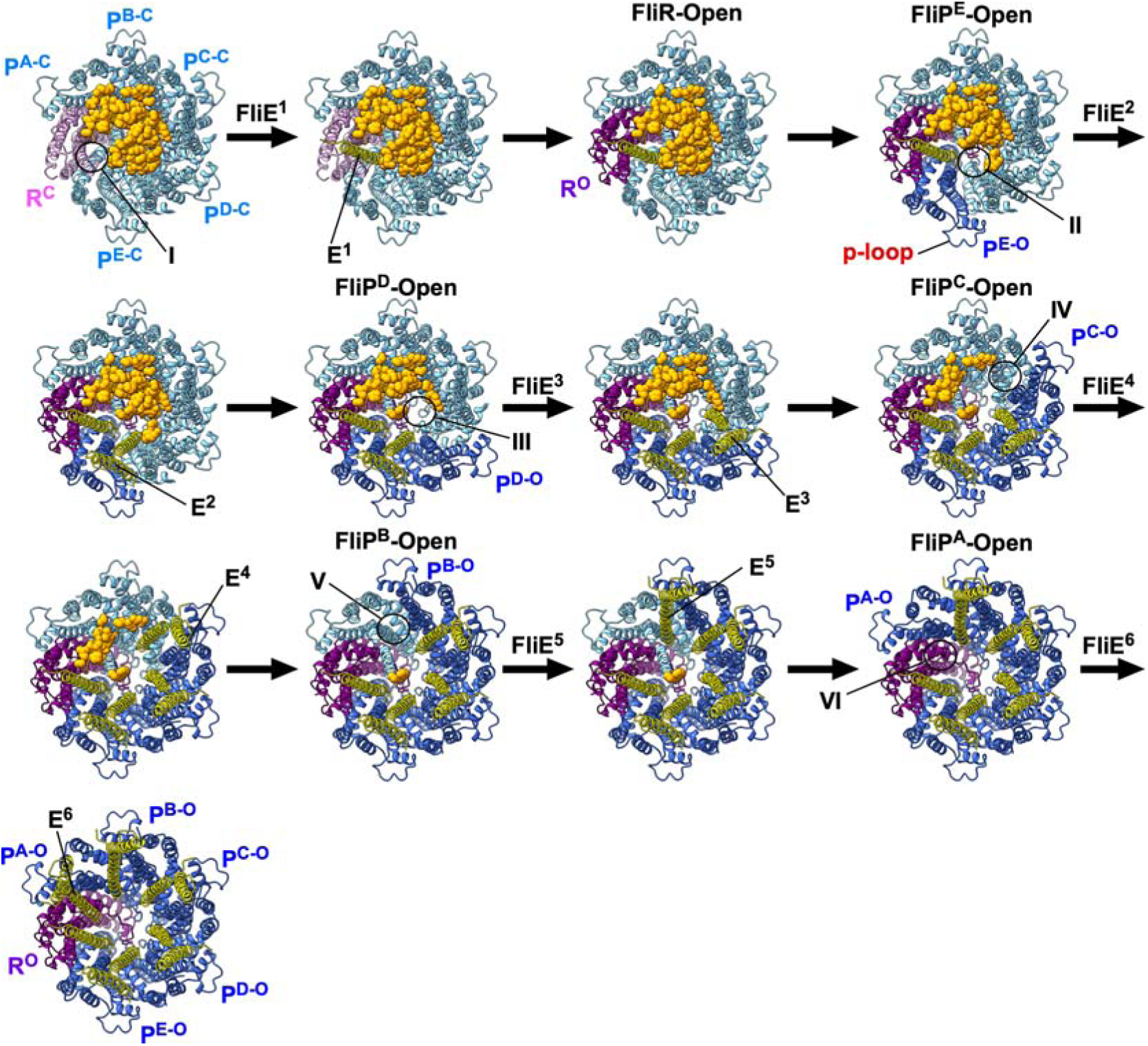
FliE assembly mechanism. The periplasmic gate of the FliPQR complex is completely closed by the β-cap while leaving a narrow pore (I), as shown in the CPK representation. The first FliE subunit (E^1^) is transported into the export channel by the fT3SS and α3 of FliE^1^ inserts into the narrow pore of the β-cap. The interaction between the closed form of FliR (R^C^) and FliE^1^ induces a conformational change of α1 of FliR, allowing this helix not only to adopt an open conformation (R^O^) but also to form the D0-like domain together with α3 of FliE^1^ (See Supplementary Fig. 10). Then, α3 of FliE^1^ associates with the closed form of the FliP-E subunit (P^E-C^), thereby inducing a conformational change of the conserved MSTF motif of FliP (See Supplementary Fig. 11). As a result, FliP^E^ adopts an open conformation (P^E-O^), creating the next insertion site (II) for the second FliE subunit (FliE^2^) within the β-cap. This open conformation is stabilized by the direct contact of the p-loop of FliP^E-O^ with the inner wall of the MS-ring. When FliE^2^ inserts into this site, α3 of FliE^2^ forms domain D0 together with α2 between FliP^E-O^ and FliP^D-C^ (P^D-C^). The interaction between FliE^2^ and FliP^D-C^ allows P^D-C^ to adopt the open conformation (P^D-O^). The FliE assembly process proceeds by repeating these steps. When the sixth FliE subunit (FliE^6^) inserts between FliP^A-O^ and FliR^O^, the periplasmic gate of the FliPQR complex is completely opened, allowing other flagellar structural subunits to diffuse through the central channel of the growing flagellum and to assemble at the distal end.

In the remaining five FliE subunits, the α2 and α3 helices form domain D0, the inner core domain of the flagellar axial structure^12,13^. Because of steric hindrances between α2 of FliE^1^ and α1 of FliR, α3 of FliE^1^ binds to α1 of FliR, thereby forming the D0-like domain (Supplementary Fig. 10a,b). Because steric hindrance also occurs between these two α-helices, α1 of FliR moves outward by removing the kink at Tyr-16 (Supplementary Fig. 10c), thereby detaching β1 of FliR from the β-cap. Consequently, FliR adopts the open form (Fig. 8, step 3), and the FliE^1^-FliR interaction is stabilized by α1 of FliP_A_ (Supplementary Fig. 11a) until the sixth FliE subunit is inserted between FliP^A^ and FliR.

The FliE^1^-FliR interaction also allows α3 of FliE^1^ to bind to α1 of FliP^E^. There is steric hindrance between these two α-helices, but Met-102 of FliE1 interacts with Leu-90 of FliP^E^ (Supplementary Fig. 11b,c) to induce a structural remodelling of the hydrophobic interaction networks surrounding the MTSF motif. As a result, both α1 and HTH_α3-α4_ of FliP^E^ move outward, not only dislodging its β-hairpin from the β-cap but also creating the insertion site for the second FliE subunit (FliE^2^) within the cap (Fig. 8, step4). The p-loop of FliP^E^ associates with the inner wall of the MS-ring, stabilizing the open conformation of FliP^E^. When FliE^2^ inserts into the second site (step5), α3 of FliE^2^ interacts with α1 of FliP^D^, which allows FliP^D^ to adopt the open conformation and creates the third FliE insertion site within the β-cap (step6). This interaction also allows α2 and α3 of FliE^2^ to form the D0 domain. By repeating this process, FliE assembly proceeds in the counterclockwise direction when viewed from the periplasmic side. When the sixth FliE subunit inserts between FliP^A^ and FliR, the periplasmic gate of the FliPQR complex is completely open.

In conclusion, the β-cap seals the periplasmic gate of the export channel to prevent premature export of flagellar proteins by the fT3SS until FliE assembles at the tip of the FliPQR complex. The β-cap also serves as a scaffold for FliE assembly. The interactions of FliE with FliP and FliR induce remodelling of the hydrophobic networks around the conserved MTSF motif, allowing the periplasmic gate, initially closed like a floral bud, to open like a blooming flower. Thus, a step-wise opening of the β-cap efficiently couples the completion of export-channel formation with the initiation of rod assembly.

## Methods

### Bacterial strains, plasmids and DNA manipulations

*Salmonella* strains and plasmids used in this study are listed in Supplementary Table 1. DNA manipulations were performed using standard protocols. Site-directed mutagenesis was carried out using Prime STAR Max Premix as described in the manufacturer’s instructions (Takara Bio). All *fliP* mutations were confirmed by DNA sequencing (Eurofins Genomics).

### Purification of the His-tagged FliPQR complex (FliPQR-His) in detergent-free solution

A 13 ml of overnight culture of *Salmonella* SJW1368 cells carrying pMKM10001 (pTrcCES3/ FliP + FliQ + FliR-His) were inoculated into a 1.3 l of fresh 2×YT [1.6% (w/v) Bacto-tryptone, 1.0% (w/v) Bacto-yeast extract, 0.5% (w/v) NaCl] containing 100 μg/ml ampicillin. The cells were grown at 30°C until the cell density had reached an OD_600_ of about 0.5–1.0, followed by the incubation at 16°C for another 24h. The cells were harvested by centrifugation (6,400 g, 10min, 4°C) and stored at −80°C. The cells were thawed, resuspended in 20 mM Tris-HCl, pH 8.0, 3 mM EDTA, and disrupted by sonication. After centrifugation (20,000 g, 15 min, 4°C) to remove cell debris, cell lysates were ultracentrifuged (110,000 g, 1 h, 4°C). The harvested membranes were solubilized in 50 mM Tris-HCl, pH 8.0, 300 mM NaCl, 5% (w/v) glycerol, 20 mM imidazole, 1% (w/v) LMNG at 4°C for 1 h and ultracentrifuged (110.000g, 1h, 4°C) to remove the insoluble membranes. Solubilized membrane proteins were loaded onto a Ni-NTA agarose column (QIAGEN) and washed extensively with 50 mM Tris-HCl, pH 8.0, 300 mM NaCl, 5% (w/v) glycerol, 20 mM imidazole, and 0.01% (w/v) LMNG. Proteins were eluted with a 100-400 mM imidazole gradient. Fractions containing FliPQR-His were concentrated, followed by size exclusion chromatography (SEC) with a Superdex 200 10/300 column (GE Healthcare) equilibrated with 20 mM Tris-HCl, pH 8.0, 300 mM NaCl, 1 mM EDTA, 5 %(w/v) glycerol, 0.005% (w/v) LMNG. Fractions containing FliPQR-His at the highest concentrations eluted from the SEC column were selected and used for reconstitution of FliPQR-His into peptidisc. The peptidisc solution was prepared from bulk lyophilized peptidisc (Peptidisc Biotech, Vancouver, BC, Canada) dissolved in 20 mM Tris-HCl, pH 7.8 to a final concentration of 5 mg/ml. Dissolved peptidisc was mixed with the solubilized FliPQR-His sample in 1:1 weight ratio and incubated at room temperature for 30 min. Then, the mixture was flowed through the Superdex 200 10/300 column equilibrated with 20mM Tris-HCl, pH 8.0, 150mM NaCl to obtain FliPQR-His in detergent-free solution.

### Sample preparation and cryoEM data collection

Fluorinated Fos-Choline was added to the purified FliPQR-His solution at a final concentration of 0, 1.0, or 2.0 mM before grid preparation. A 2.7 μl aliquot of the sample solution (2.0 mg/ml) were applied onto a glow-discharged holey carbon-coated grid (Quantifoil 200mesh, Cu R1.2/1.3). The grid was blotted by a filter paper at 4°C for 3 sec and quickly frozen in liquid ethane using a Vitrobot Mark IV system (Thermo Fisher Scientific, 4°C and 100% humidity).

The grids were inserted into a CRYO ARM 300 transmission electron microscopy (JEOL Ltd. Japan) equipped with a cold field-emission electron gun operated at 300□kV and an Ω-type energy filter with a 20 eV slit width. CryoEM images were recorded with a K3 direct electron detector camera (Gatan, USA) at a nominal magnification of ×60,000, corresponding to an image pixel size of 1.0 Å, using SerialEM^46^. The holes were detected using YoneoLocr^47^. Movie frames were recorded in CDS counting mode with a total exposure time of 3 sec and a total dose of ∼40 electrons Å^−2^. Each movie was fractionated into 40 frames. In total, 13,230 movies were collected.

### CryoEM image processing

Single particle analysis was performed using RELION 3.1^48^. Image processing procedure is described in Supplementary Fig. 1b. After performing motion corrections to align all micrographs, followed by the estimation of parameters of the contrast transfer fraction (CTF), particle images were automatically selected via LoG auto-picking, and the selected particles were extracted into a box of 256 × 256 pixels (1,505,398 particles). Particle images from good 2D class average images were selected for the initial 3D model. In total, 164,019 particles were subjected to 3D classification with C1 symmetry. After 3D refinement for each of this good one class, postprocessing yielded the 3D map at a resolution of 3.0 Å (109,333 particles) according to 0.143 criterion of the Fourier shell correlation (FSC). The cryoEM density map was deposited into Electron Microscopy Data Bank with an accession code EMD-61993.

### Model building and refinement of the FliPQR complex

The atomic model of the FliPQR complex was constructed using Coot^49^. PHENIX was used for real-space refinement based on the cryoEM map^50^. Summary of model refinement and statistics are described in Table 1. The atomic coordinates have been deposited in the Protein Data Bank with an accession code 9K29.

### Motility assays in soft agar

Fresh colonies were inoculated into soft agar plates [1% (w/v) triptone, 0.5% (w/v) NaCl, 0.35% Bacto agar] containing 100 μg/ml ampicillin and incubated at 30°C. At least five independent measurements were performed.

### Secretion assay

A 100 μl of the overnight culture of *Salmonella* cells was inoculated into a 5 ml of fresh L-broth [1% (w/v) triptone, 0.5% (w/v) yeast extract, 0.5% (w/v) NaCl] containing 100 μg/ml ampicillin and incubated at 30 °C with shaking until the cell density had reached an OD_600_ of ca. 1.4–1.6. Cultures were centrifuged to obtain cell pellets and culture supernatants, separately. The cell pellets were resuspended in sodium dodecyl sulfate (SDS)-loading buffer solution [62.5 mM Tris-HCl, pH 6.8, 2% (w/v) SDS, 10% (w/v) glycerol, 0.001% (w/v) bromophenol blue] containing 1 μl of 2-mercaptoethanol. Proteins in each culture supernatant were precipitated by 10% trichloroacetic acid and suspended in a Tris/SDS loading buffer (one volume of 1 M Tris, nine volumes of 1 X SDS-loading buffer solution) containing 1 μl of 2-mercaptoethanol. Both whole cellular proteins and culture supernatants were normalized to a cell density of each culture to give a constant cell number. After boiling at 95°C for 3 min, the samples were separated by sodium dodecyl sulfate-polyacrylamide gel electrophoresis (SDS-PAGE) and transferred to a nitrocellulose membrane (Bio-Rad) using a transblotting apparatus (Hoefer). Then, immunoblotting with polyclonal anti-FlgD or anti-FliP antibody as the primary antibody and anti-rabbit IgG, HRP-linked whole Ab Donkey (GE Healthcare) as the secondary antibody was carried out using iBand Flex Western Device as described in the manufacturer’s instructions (Thermo Fisher Scientific). Detection was performed with Amersham ECL Prime western blotting detection reagent (Cytiva). Chemiluminescence signals were captured by a Luminoimage analyzer LAS-3000 (GE Healthcare). Bands of prestained protein molecular weight markers (Bio-Rad) transferred to each membrane were also photographed with the LAS-3000 under brightfield illumination and combined with each immunoblot image to identify the band of interest. All image data were processed with Photoshop software (Adobe). At least three independent experiments were performed.

### Multiple sequence alignment

Multiple sequence alignment was carried out using Clustal Omega (https://www.ebi.ac.uk/jdispatcher/msa/clustalo).

### Data availability

The cryoEM map and atomic model of the *Salmonella* FliPQR complex reconstituted in a peptidisc have been deposited in the Electron Microscopy Data Bank under an accession code EMD-61993 and the Protein Data Bank under an accession code 9K29. All data generated during this study are included in this published article, and its Supplementary Information. Strains, plasmids, polyclonal antibodies, and all other data are available from the corresponding author upon request.

## Acknowledgements

We acknowledge Michael D. Manson for critical reading of the manuscript and helpful comments, Kelly T. Hughes for his kind gift of the *Salmonella* Δ*fliP* mutants and Yasuyo Abe and Yoshie Kushima for technical assistance. This work was supported in part by JSPS KAKENHI Grant Numbers JP20K15749 and JP22K06162 (to M.K.) and JP19H03182, JP22H02573, and JP22K19274 (to T.Minamino) and MEXT KAKENHI Grant Numbers JP20H05532, and JP22H04844 (to T.Minamino). This work has also been supported by Platform Project for Supporting Drug Discovery and Life Science Research (BINDS) from AMED under Grant Number JP19am0101117 and JP21am0101117 (to K.N.), by the Cyclic Innovation for Clinical Empowerment (CiCLE) from AMED under Grant Number JP17pc0101020 (to K.N.), and by JEOL YOKOGUSHI Research Alliance Laboratories of Osaka University (to K.N.).

## Author Contributions

K.N. and T.Minamino conceived and designed research; M.K. prepared samples for cyroEM; M.K., T.Miyata and F.M. collected and analysed cryoEM image data; M.K. and K.I. built atomic models; M.K. and T.Minamino performed genetic, biochemical, and physiological experiments; M.K., K.N. and T.Minamino wrote the paper based on discussion with other authors.

