## Supplementary Information for "Structural basis of flagellar rod assembly on the FliPQR protein-export channel"

### Supplementary Result 1

**Mutational analysis of well-conserved residues forming hydrophobic side-chain interaction networks surrounding the MTSF motif.** The hydrophobic parts of the side-chains of Met-61, Thr-62, Ser-63, and Phe-64 form hydrophobic interaction networks with those of the surrounding residues, thereby stabilizing a closed conformation of the MTSF motif (Supplementary Figs. 6a and 7). This raises the possibility that a conformational change in the MTSF motif occurs through the reorganization of the hydrophobic interaction networks surrounding the MTSF motif. To clarify this possibility, we first chose five residues, Leu-90, Leu-92, Leu-96, Thr-97, and Phe-98 and replaced each with alanine. These alanine substitutions did not affect the cellular level of FliP (Supplementary Fig. 6b). The L90A substitution slightly reduced motility in soft agar, and the L92A substitution inhibited motility (Fig. 6a and Supplementary Fig. 6c). However, the remaining three alanine substitutions did not affect the motility. These results indicate that Leu-92 is critical for FliP function.

The conserved Thr-97 residue establishes intramolecular hydrophobic contacts with Thr-62 and Phe-64 in the MTSF motif. The conserved Leu-96 residue forms a hydrophobic core along with Phe-71, Leu-92, Met-236, and Leu-239. The conserved Phe-98 residue engages in an intermolecular interaction with the hydrophobic part of the side chain of Arg-168 of its nearest FliP subunit (Supplementary Figs. 6a and 7). Therefore, we next constructed four *fliP* mutants, *fliP*(L96A/T97A), *fliP*(T97A/F98A), *fliP*(L96A/F98A), and *fliP*(L96A/T97A/F98A). These substitutions did not largely affect the cellular level of FliP. The L96A/T97A double substitution led to a reduction in motility by approximately 2-fold, and the L96A/T97A/F98A triple substitution resulted in a strong inhibition of motility (Supplementary Fig. 6c, right panel). In contrast, the T97A/F98A and L96A/F98A double substitution had no impact on motility. These observations suggest that a conformational change of the MTSF motif of FliP occurs through proper reorganization of the hydrophobic interaction networks surrounding the MTSF motif, allowing helix α1 of FliP to switch from the closed conformation to the open conformation.

### Supplementary Result 2

**Mutational analysis of well-conserved residues in helix α1 of FliP.** In addition to the β-cap, intermolecular hydrophobic interactions between the α1 helices of the five FliP subunits firmly close the periplasmic gate of the export channel in the closed structure of the FliPQR complex. We found that the *fliP*(L45Q) mutation in α1 of FliP destabilizes the closed conformation of FliP(L92A), thereby enabling FliP(L92A) to adopt the open conformation (Fig. 6). This suggests that the hydrophobicity of Leu-45 is functionally important. If this is true, then replacing this residue with alanine will not affect FliP function. To clarify this hypothesis, we chose five conserved residues, Leu-45, Phe-47, Leu-51, Thr-52, and Phe-53 and replaced each with alanine (Supplementary Fig. 8a). As expected, these alanine substitutions did not affect protein stability (Supplementary Fig. 8b) or motility (Supplementary Fig. 8c). These conserved residues play a crucial role in establishing the hydrophobic core between the five $\alpha1_a$ helices, thereby closing the export channel in the 9K29 structure (Supplementary Fig. 9, upper panels). In contrast, these residues are also involved in the interaction between FliP and FliE (Supplementary Fig. 5), thereby opening the export channel in the 8WKK structure (Supplementary Fig. 9, lower panels). Therefore, we suggest that the hydrophobic nature of α1 of FliP plays a critical role in the proper closing and opening of the periplasmic gate of the export channel.

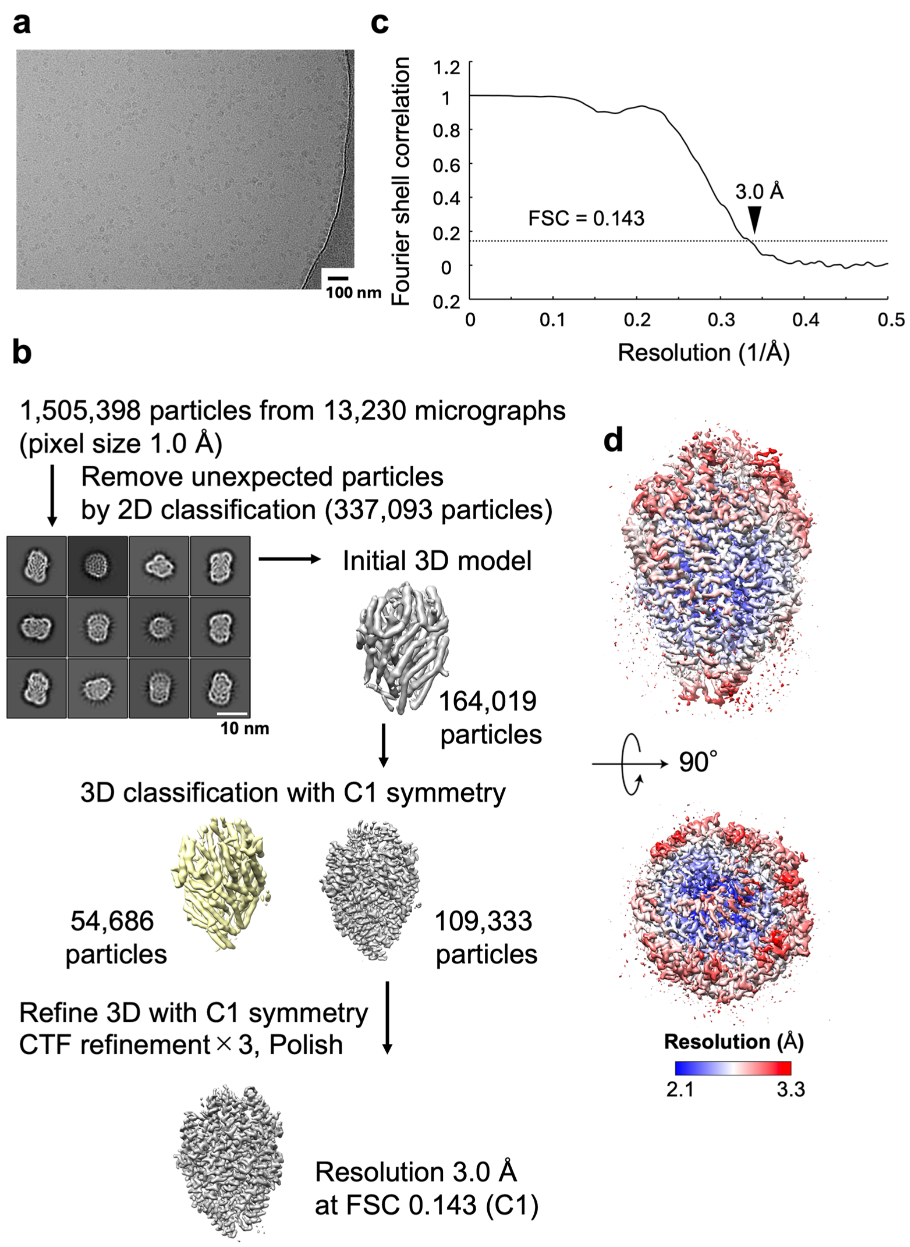

**Supplementary Fig. 1. CryoEM single particle 3D image analysis of the FliPQR complex reconstituted in a peptidisc. (a)** Representative cryoEM image of the FliPQR complex reconstituted in the peptidisc. **(b)** Work process of cryoEM single particle 3D reconstruction. **(c)** Fourier shell correlation (FSC) curve of the final density map of the FliPQR complex (EMDB ID: EMD-61993. **(d)** Local resolution of the final density map of the FliPQR complex is colored from blue (2.1 Å) to red (3.3 Å).

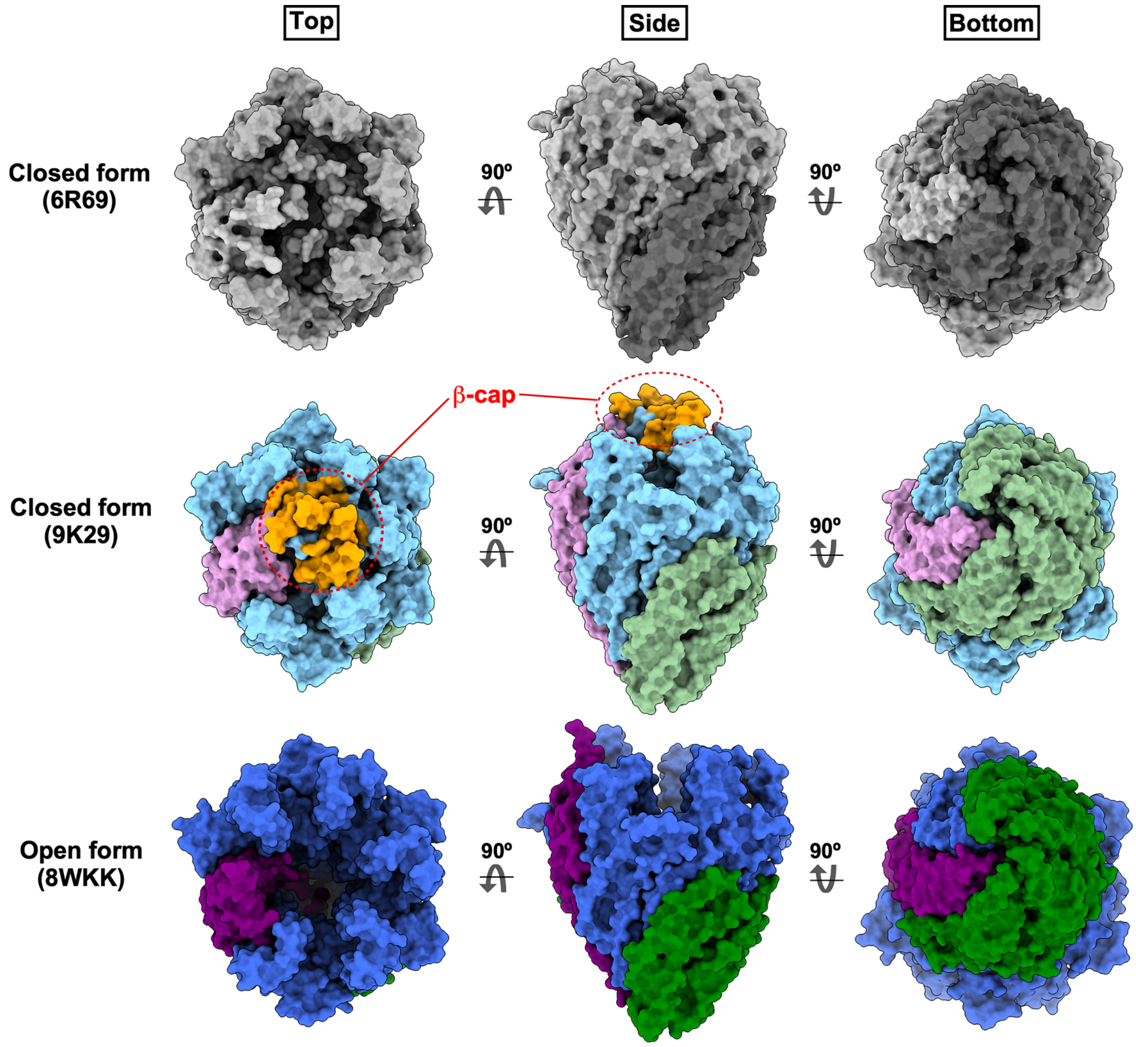

**Supplementary Fig. 2. Surface maps of three different cryoEM structures of FliPQR complex.** In the 6R69 (top panels) and 9K29 (middle panels) structures, the periplasmic gate of the FliPQR complex is closed by intermolecular hydrophobic interactions between the N-terminal α1 helices of the five FliP and one FliR subunits but is open in the 8WKK structure (lower panels). The periplasmic gate is completely closed by the β-cap in the 9K29 structure. In contrast, the β-cap is missing in the 6R69 and 8WKK structures. The cytoplasmic gate is closed in these three different structures.

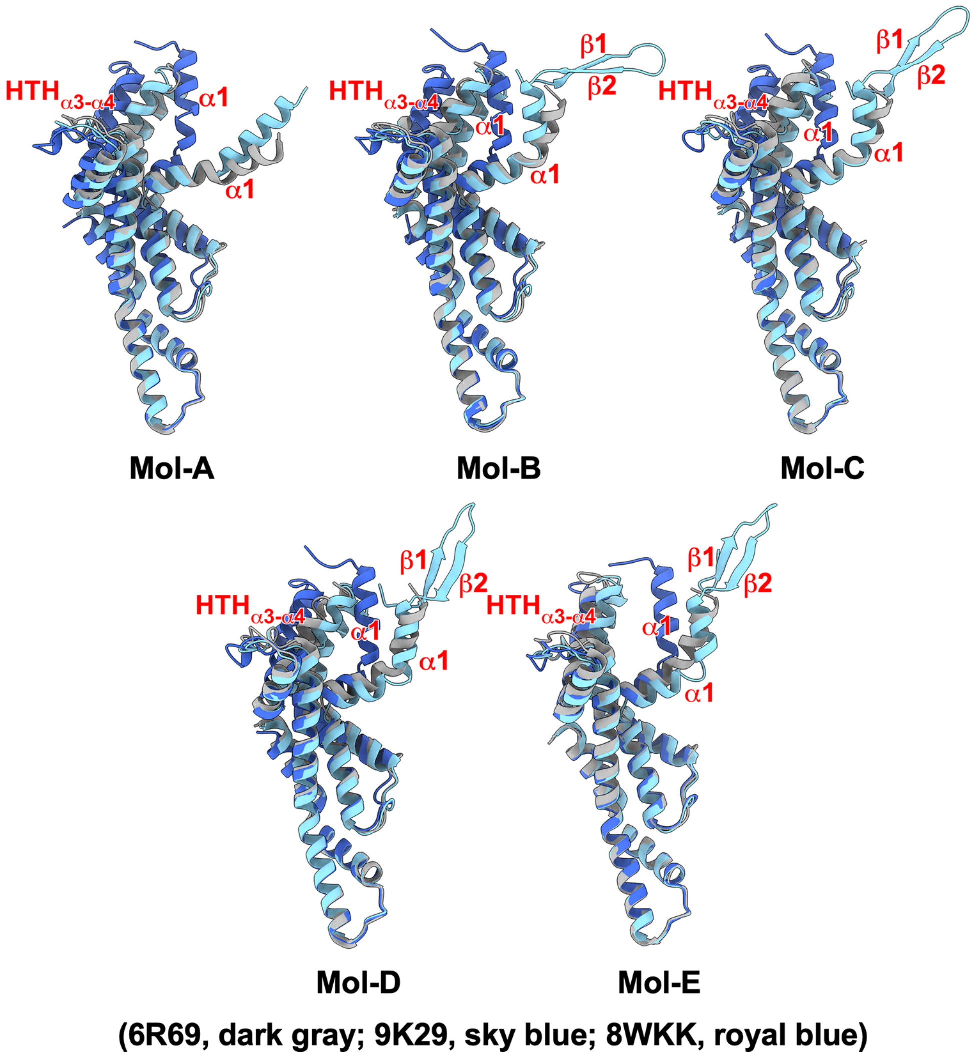

**Supplementary Fig. 3. Superpositions of each FliP subunit obtained in this study (PDB ID: 9K29) with the equivalent coordinates of the 6R69 and 8WKK structures.** Each FliP subunit of the 6R69 (dark gray) and 8WKK (royal blue) structures were superimposed on the corresponding FliP subunit of the 9K29 structure (sky blue).

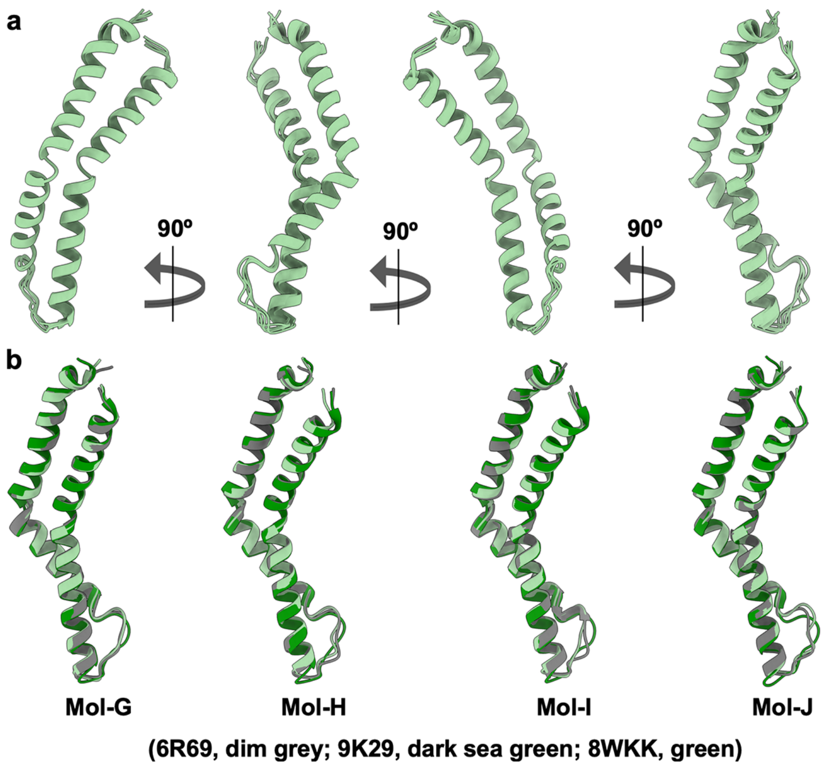

**Supplementary Fig. 4. Structural comparisons of the four FliQ subunits in the FliPQR complex. (a)** Structural comparisons of the four FliQ subunits in the 9K29 structure. They are all nearly identical to one another. **(b)** Superposition of each FliQ subunit obtained in this study (PDB ID: 9K29) (dark sea green) with the equivalent coordinates of the 6R69 (dim grey) and 8WKK (green) structures. Each FliQ subunit of the 6R69 and 8WKK structures were superimposed on the corresponding FliQ subunit of the 9K29 structure.

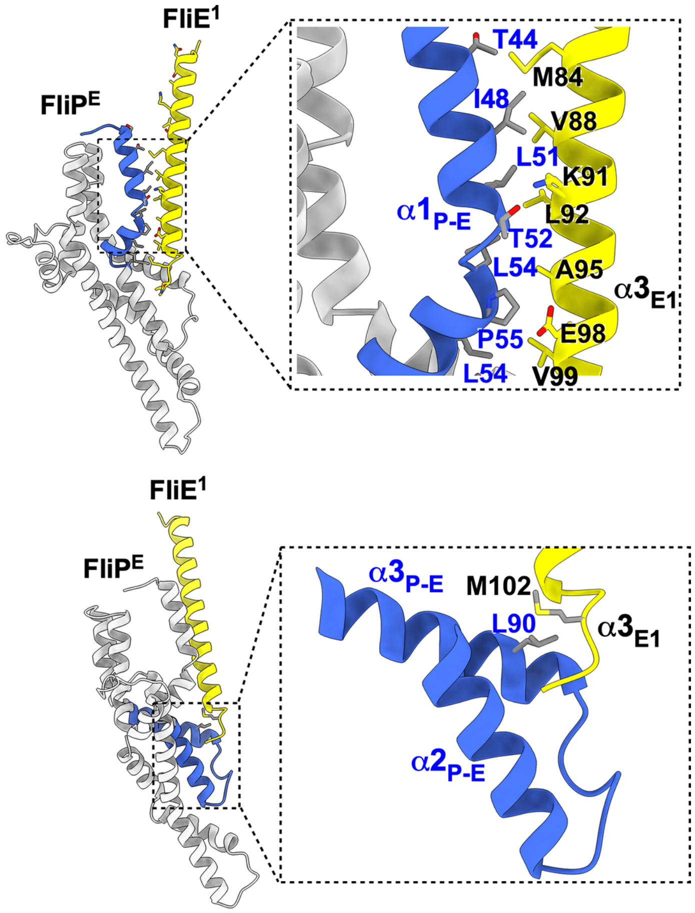

**Supplementary Fig. 5. Interaction between FliP and FliE in the 8WKK structure.** Helix α3 of the first FliE subunit (FliE$^1$) firmly associates with helix α1 of the FliP-E subunit (FliP$^E$). The C-terminal end of FliE$^1$ also binds to the helix-turn-helix structure formed by helices α2 and α3 of FliP$^E$. Leu-90 of FliP$^E$ makes hydrophobic contact with Met-102 of FliE$^1$

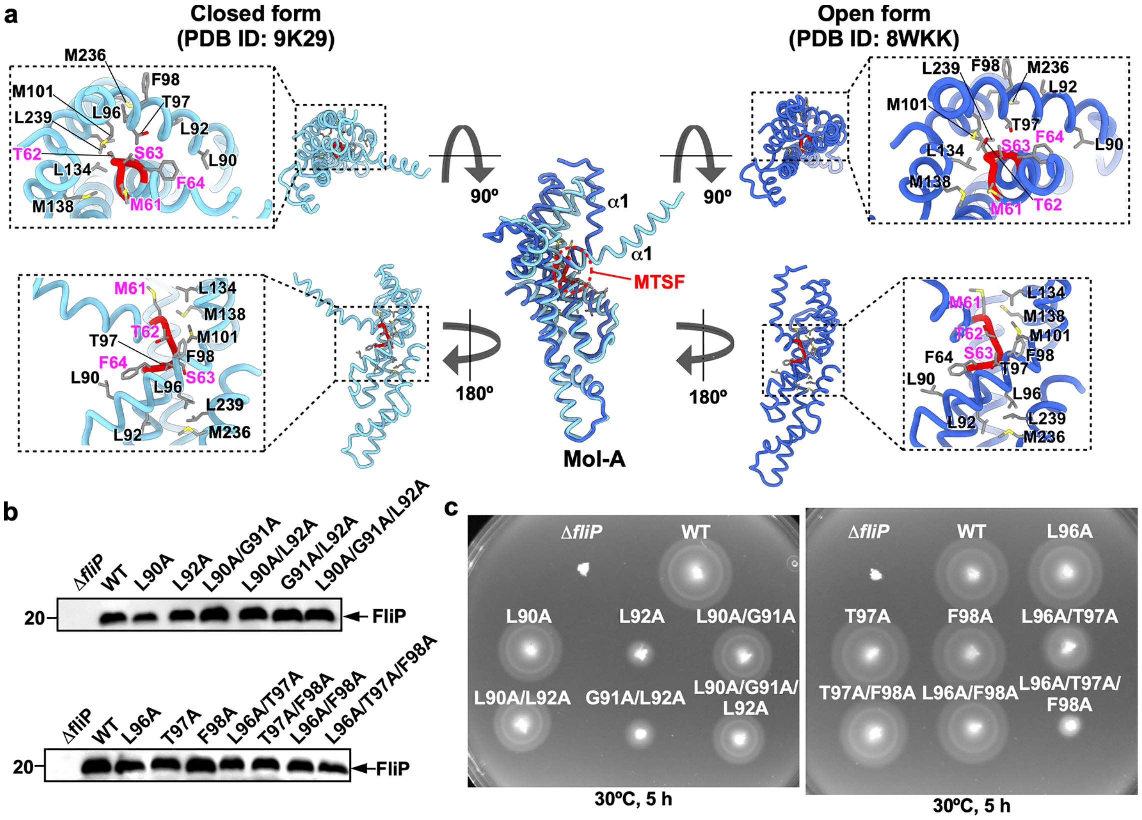

**Supplementary Fig. 6. Mutational analysis of conserved residues forming hydrophobic side-chain interaction networks around the MTSF motif. (a)** Hydrophobic interaction networks surrounding the MTSF motif (red) in the closed and open forms of FliP. **(b)** Immunoblotting, using polyclonal anti-FliP antibody, of whole cell lysates prepared form a *Salmonella fliP* null mutant harboring pTrc99AFF4 (Δ*fliF*), pKY69 (WT), pMKM69(L90A) (indicated as L90A), pMKM69(L92A) (L92A), pMKM69(L90A/G91A) (L90A/G91A), pMKM69(L90A/L92A) (L90A/L92A), pMKM69(G91A/L92A) (G91A/L92A), pMKM69(L90A/G91A/L92A) (L90A/G91A/L92A), pMKM69(L96A) (L96A), pMKM69(T97A) (T97A), pMKM69(F98A) (F98A), pMKM69(L96A/T97A) (L96A/T97A), pMKM69(T97A/F98A) (T97A/F98A), pMKM69(L96A/F98A) (L96A/F98A), or pMKM69(L96A/T97A/F98A) (L96A/T97A/F98A) The position of the marker with a molecular wight of 20 kDa is shown on the left. At least three independent assays were performed. **(c)** Motility of the above transformants in soft agar. The plate was incubated at 30ºC for 5 hours. At least seven independent assays were carried out.

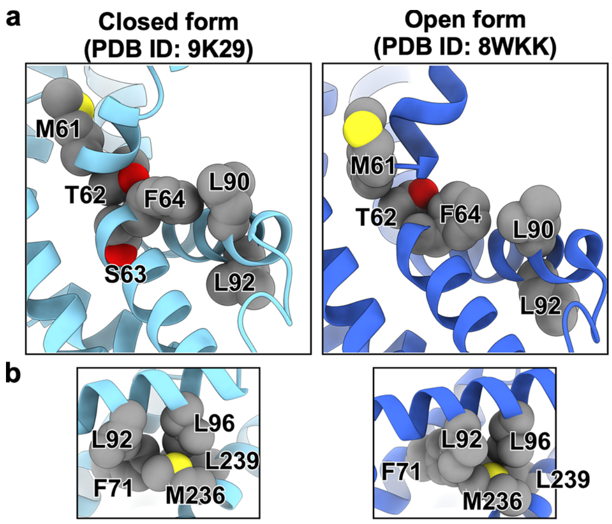

**Supplementary Fig. 7. The interaction between FliE and FliP causes a conformational transition from the closed to the open state of FliP. (a)** Conformational change of the MTSF motif by restructuring the hydrophobic interaction network around the MTSF motif. **(b)** Remodeling of the hydrophobic interaction network formed by Phe-71, Leu-92, Leu-96, Met-236, and Leu-239.

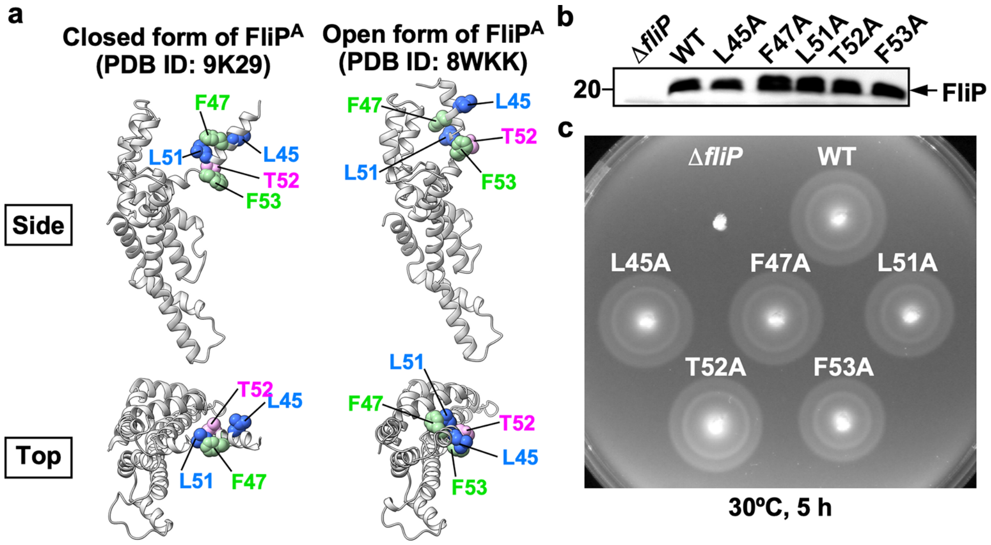

**Supplementary Fig. 8. Mutational analysis of conserved Leu-45, Phe-47, Leu-51, Thr-52, and Phe-53 residues of FliP. (a)** Location of Leu-45, Phe-47, Leu-51, Thr-52, and Phe-53 in the FliP structure. **(b)** Immunoblotting, using polyclonal anti-FliP antibody, of whole cell lysates prepared form a *Salmonella* Δ*fliP* mutant transformed with pTrc99FF4 (Δ*fliP*), pKY69 (WT), pMKM69(L45A) (L45A), pMKM69(F47A) (F47A), pMKM69(L51A) (L51A), pMKM69(T52A) (T52A) or pMKM69(F53A) (F53A). The position of the marker with a molecular wight of 20 kDa is shown on the left. At least three independent assays were performed. **(c)** Motility of the above transformants. The plate was incubated for at 30ºC for 5 hours. At least seven independent assays were carried out.

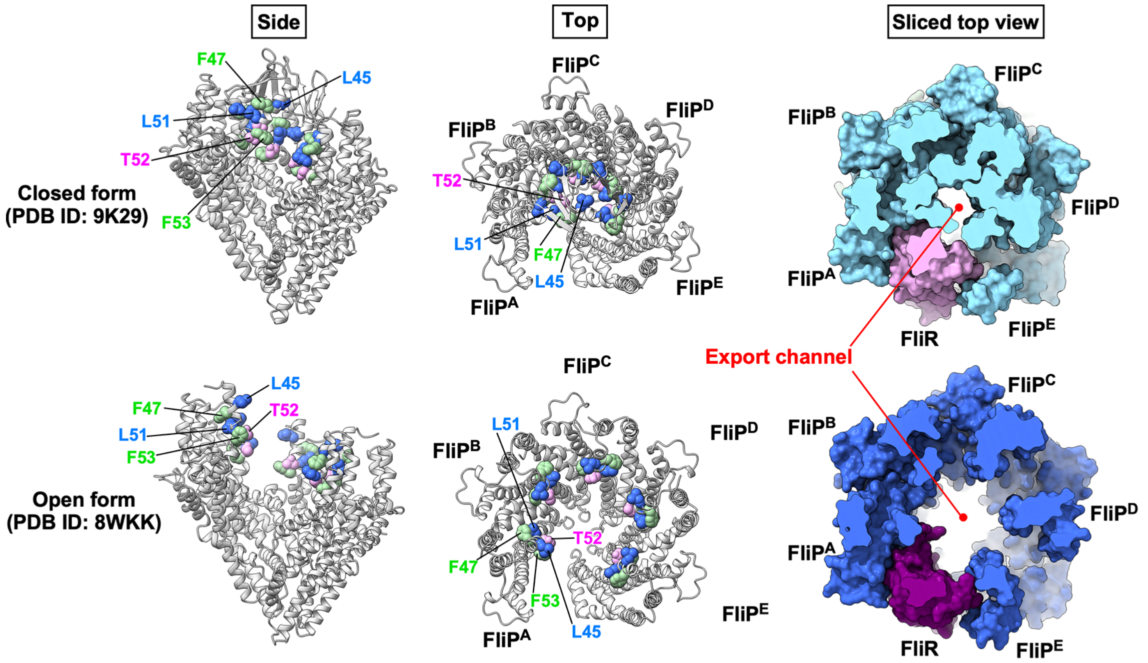

**Supplementary Fig. 9. The protein export channel is closed by a hydrophobic core formed by hydrophobic residues in the N-terminal α1 helices of the five FliP subunits in the 8WKK structure.** Leu-45, Phe-47, Leu-51, Thr-52, and Phe-53 are well-conserved among FliP homologues. These residues are in α1 of FliP and form a hydrophobic core in the closed FliPQR complex (PDB ID: 9K29). They are also involved in the interaction with FliE in the 8WKK structure. The right panels show the horizontally sliced top views of the surface maps.

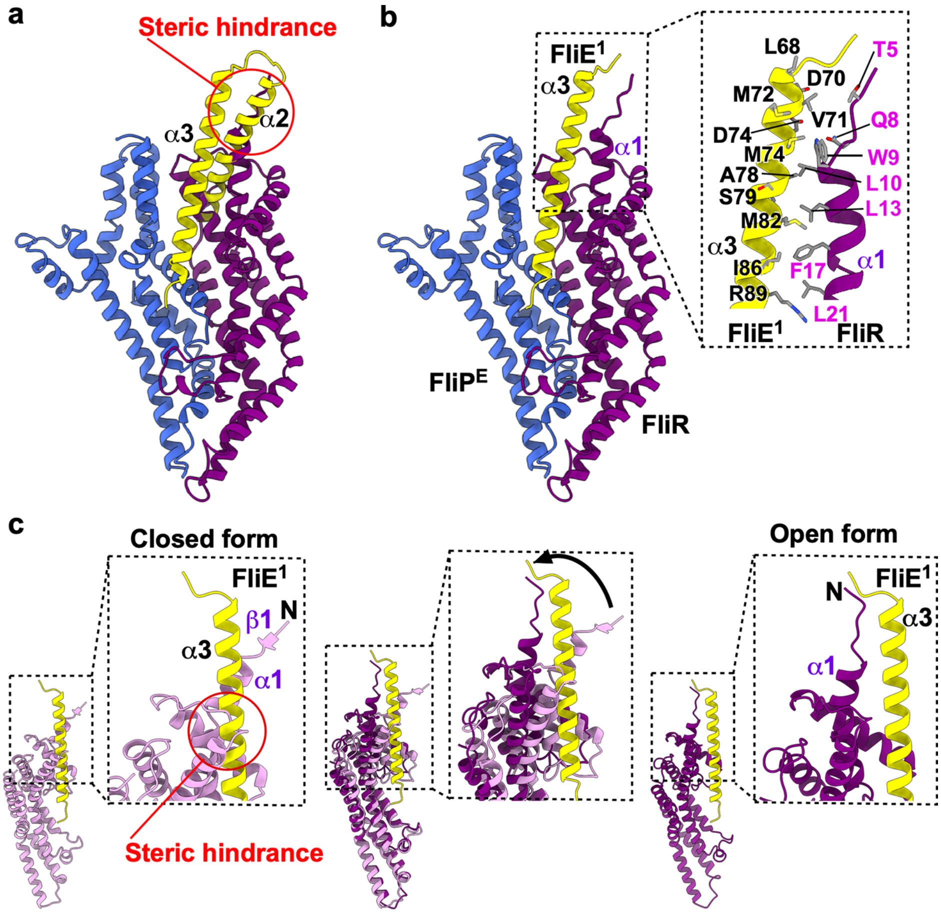

**Supplementary Fig. 10. Interaction between the first FliE subunit and FliR. (a)** Interaction between domain D0 of FliE and FliR. FliE (yellow) contains three α-helices with the α2 and α3 helices forming the D0 domain. Domain D0 of FliE is superimposed on α3 of the first FliE subunit ($FliE^1$) in the 8WKK structure. Because α2 of the D0 domain collides with α1 of FliR at the position indicated by the red circle, α2 of $FliE^1$ cannot form domain D0 along with α3. Consequently, α2 of $FliE^1$ is invisible in the 8WKK structure. **(b)** Interaction between α3 of $FliE^1$ and α1 of the open form of FliR (purple) (PDB ID: 8WKK). When $FliE^1$ inserts between the FliR and $FliP^E$ subunits, its C-terminal α3-helix makes hydrophobic contacts with α1 of FliR to form the D0-like domain. **(c)** Interaction between $FliE^1$ and the closed form of FliR. The equivalent coordinates of the 9K29 (plum) and 8WKK (purple) structures are superimposed. In the closed structure of FliR, α3 of FliE collides with α1 of FliR at the position indicated by the red circle. This collision allows α1 of FliR to move outward, thereby dislodging the β-strand (β1) from the β-cap. Consequently, α3 of FliE forms the D0-like domain together with α1 of the open form of FliR.

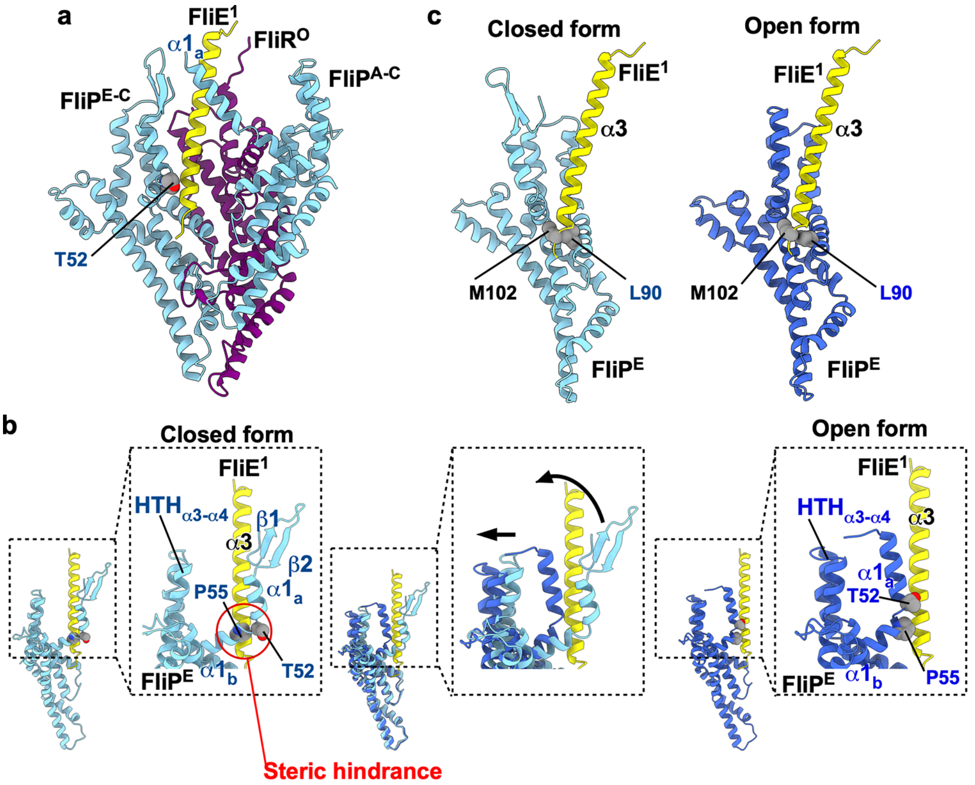

**Supplementary Fig. 11. Interaction between the first FliE subunit and the closed form of the FliP-E subunit. (a)** The equivalent coordinates of the 9K29 and 8WKK structures are superimposed. Helix $\alpha 1_a$ of the closed FliP-A subunit ($FliP^{A\text{-}C}$) supports efficient insertion of the first FliE subunit ($FliE^1$) between the open form of FliR ($FliR^O$) and the closed form of FliP-E ($FliP^{E\text{-}C}$). **(b)** Interaction between Met-102 of $FliE^1$ and Leu-90 of FliP. Met-102 of $FliE^1$ makes hydrophobic contact with Leu-90 of FliP-E in both closed and open conformations. This interaction is presumed to reconstitute a hydrophobic side-chain interaction network surrounding the MTSF motif, causing a conformational change of the MTSF motif. **(c)** Interaction between α3 of $FliE^1$ and α1 of $FliP^E$. When α3 of $FliE^1$ is proximal to $\alpha 1_a$ of the closed form of $FliP^E$, the C-terminal portion of α3 of $FliE^1$ collides with Thr-52 and Pro-55 of $FliP^E$ at the position indicated by the red circle. The conformational change of the MTSF motif leads to the outward movement of α1 and the helix-turn-helix structure formed by the α3 and α4 helices ($HTH_{\alpha 3\text{-}\alpha 4}$), resulting in the dislodgment of the β-hairpin of $FliP^E$ from the β-cap. As a result, FliP adopts the open conformation and firmly associates with α3 of $FliE^1$.

**Supplementary Table 1. Strains and plasmids used in this study**

| Strains and Plasmids | Relevant characteristics | Source or reference |
|---|---|---|
| ***Salmonella*** | | |
| SJW1103 | Wild-type for motility and chemotaxis | 1 |
| SJW1368 | Δ(*cheW-flhD*) | 2 |
| TH10549 | Δ*fliP* | K. T. Hughes |
| **Plasmid** | | |
| pTrc99AFF4 | Modified pTrc99A | 3 |
| pTrcES | Modified pTrc99A | This study |
| pKY69 | pTrc99AFF4/ His-FliP | 4 |
| pMKM10001 | pTrcES/ FliP + FliQ + FliR-His | This study |
| pMKM69(ΔMTSF) | pTrc99AFF4/ His-FliP(ΔMTSF) | This study |
| pMKM69(T62A/S63A) | pTrc99AFF4/ His-FliP(T62A/S63A) | This study |
| pMKM69(T62G/S63G) | pTrc99AFF4/ His-FliP(T62G/S63G) | This study |
| pMKM69(L45A) | pTrc99AFF4/ His-FliP(L45A) | This study |
| pMKM69(F47A) | pTrc99AFF4/ His-FliP(F47A) | This study |
| pMKM69(L51A) | pTrc99AFF4/ His-FliP(L51A) | This study |
| pMKM69(T52A) | pTrc99AFF4/ His-FliP(T52A) | This study |
| pMKM69(F53A) | pTrc99AFF4/ His-FliP(F53A) | This study |
| pMKM69(L90A) | pTrc99AFF4/ His-FliP(L90A) | This study |
| pMKM69(L92A) | pTrc99AFF4/ His-FliP(L92A) | This study |
| pMKM69(L90A/G91A) | pTrc99AFF4/ His-FliP(L90A/G91A) | This study |
| pMKM69(L90A/L92A) | pTrc99AFF4/ His-FliP(L90A/L92A) | This study |
| pMKM69(G91A/L92A) | pTrc99AFF4/ His-FliP(G91A/L92A) | This study |
| pMKM69(L90A/G91A/L92A) | pTrc99AFF4/ His-FliP(L90A/G91A/L92A) | This study |
| pMKM69(L92A)-SP1 | pTrc99AFF4/ His-FliP(P30L/L92A) | This study |
| pMKM69(L92A)-SP2 | pTrc99AFF4/ His-FliP(L45Q/L92A) | This study |
| pMKM69(L92A)-SP3 | pTrc99AFF4/ His-FliP(L92A/R168C) | This study |
| pMKM69(L96A) | pTrc99AFF4/ His-FliP(L96A) | This study |
| pMKM69(T97A) | pTrc99AFF4/ His-FliP(T97A) | This study |
| pMKM69(F98A) | pTrc99AFF4/ His-FliP(F98A) | This study |
| pMKM69(L96A/T97A) | pTrc99AFF4/ His-FliP(L96A/T97A) | This study |
| pMKM69(T97A/F98A) | pTrc99AFF4/ His-FliP(T97A/F98A) | This study |
| pMKM69(L96A/F98A) | pTrc99AFF4/ His-FliP(L96A/F98A) | This study |
| pMKM69(L96A/T97A/F98A) | pTrc99AFF4/ His-FliP(L96A/T97A/F98A) | This study |
| pMKM69(L96A/T97A/F98A)-SP5 | pTrc99AFF4/ His-FliP(L96E/T97A/F98A) | This study |
| pMKM69(Δ155-164) | pTrc99AFF4/ His-FliP(Δ155-164) | This study |
| pMKM69(Δ155-165) | pTrc99AFF4/ His-FliP(Δ155-165) | This study |
| pMKM69(Δ156-163) | pTrc99AFF4/ His-FliP(Δ156-163) | This study |
| pMKM69(Δ157-162) | pTrc99AFF4/ His-FliP(Δ157-162) | This study |
